# Effects of FLOWERING LOCUS T on FD during the transition to flowering at the shoot apical meristem of *Arabidopsis thaliana*

**DOI:** 10.1101/483925

**Authors:** Silvio Collani, Manuela Neumann, Levi Yant, Markus Schmid

## Abstract

The transition to flowering is a crucial step in the plant life cycle that is controlled by multiple endogenous and environmental cues, including hormones, sugars, temperature, and photoperiod. Permissive photoperiod induces *FLOWERING LOCUS T* (*FT*) in the phloem companion cells of leaves. The FT protein then acts as a florigen that is transported to the shoot apical meristem (SAM) where it physically interacts with the bZIP transcription factor FD and 14-3-3 proteins. However, despite the importance of FD for promoting flowering, its direct transcriptional targets are largely unknown. Here we combined ChIP-seq and RNA-seq to identify targets of FD at the genome-wide scale and assess the contribution of FT to binding DNA. We further investigated the ability of FD to form protein complexes with FT and TFL1 through the interaction with 14-3-3 proteins. Importantly, we observe direct binding of FD to targets involved in several aspects of the plant development not directly related to the regulation of flowering time. Our results confirm FD as central regulator of the floral transition at the shoot meristem and provides evidence for crosstalk between the regulation of flowering and other signaling pathways.

**Material Distribution:** The author responsible for distribution of materials integral to the findings presented in this article in accordance with the policy described in the Instructions for Authors (www.cell.com/molecular-plant/authors): Markus Schmid.

**Contact Information:** Umeå Plant Science Centre (UPSC), Dept. of Plant Physiology Umeå University, SE-901 87 Umeå, SWEDEN

## INTRODUCTION

The floral transition represents a crucial checkpoint in the plant life cycle at which the shoot apical meristem (SAM) stops producing only leaves and begins producing reproductive organs. As the commitment to this developmental phase transition is usually irreversible for a given meristem, plants have evolved several pathways to integrate environmental and endogenous stimuli to ensure flowering is induced at the correct time. A rich literature has identified hormones, sugars, temperature, and day length (photoperiod) as main factors in flowering time regulation (reviewed in Romera-Branchat et al., 2014; Song et al., 2015; Srikanth and Schmid, 2011). Photoperiod in particular has been shown to regulate flowering time in many plant species and, depending on the light requirements, short day (SD), long day (LD) and day-neutral plants have been distinguished. In *Arabidopsis thaliana*, LD promotes flowering but plants will eventually flower even under non-inductive SD.

It has long been known that in day-length responsive species, inductive photoperiod is mainly perceived in leaves where it results in the formation of a long-distance signal, or florigen, that moves to the SAM to induce the transition to flowering (An et al., 2004; Corbesier et al., 2007; Mathieu et al., 2007). The molecular nature of florigen has eluded identification for the better part of a century. However, recently *FLOWERING LOCUS T* (*FT*) and related genes, which encode for phosphatidylethanolamine-binding proteins (PEBP), have been identified as evolutionary conserved candidates (Corbesier et al., 2007; Mathieu et al., 2007). Under inductive photoperiod, *FT* is expressed in leaf phloem companion cells (PCC) and there is good evidence that the FT protein is loaded into the phloem sieve elements and transported to the SAM (reviewed in (Song et al., 2015; Srikanth and Schmid, 2011)). At the SAM, FT interacts with FD and 14-3-3 proteins and the resulting flowering-activation complex (FAC) is thought to control the correct expression of flowering time and floral homeotic genes to promote the transition of the vegetative meristem into a reproductive inflorescence meristem (Abe et al., 2005; Taoka et al., 2011; Wigge et al., 2005).

FD belongs to the group A of the bZIP transcription factor (TF) family (Jakoby et al., 2002) and is mainly expressed at the SAM (Abe et al., 2005; Schmid et al., 2005; Wigge et al., 2005). It has been proposed that, in order to interact with FT and 14-3-3 proteins, FD must be phosphorylated at threonine 282 (T282) (Abe et al., 2005; Taoka et al., 2011; Wigge et al., 2005). Recently, two calcium-dependent kinases expressed at the SAM, CPK6 and CPK33, have been shown to phosphorylate FD (Kawamoto et al., 2015). FD interacts not only with FT but also with other members of the PEBP protein family. Interestingly, some of the six PEBP proteins encoded in the *A. thaliana* genome regulate flowering in opposition. FT and its paralog TWIN SISTER OF FT (TSF) promote flowering. Mutations in *tsf* enhance the late flowering phenotype of *ft* in LD but in addition *TSF* also has distinct roles in SD (Yamaguchi et al., 2005). Other members of the PEBP protein family, most prominently TERMINAL FLOWER 1 (TFL1), oppose the flower-promoting function of FT and TSF, and repress flowering. The Arabidopsis ortholog of CENTRORADIALIS (ATC) has been shown to act as a SD-induced floral inhibitor that is expressed mostly in the vasculature but was undetectable at the SAM. Furthermore, ATC has been suggested to move over long distances and can interact with FD to inhibit *APETALA1* (*AP1*) expression. ATC has thus been proposed to antagonize the flower-promoting effect of FT (Huang et al., 2012). Similarly, orthologs of ATC in rice (RCNs) have been recently showed to antagonize with FT-like protein (Kaneko-Suzuki et al., 2018). Finally, BROTHER OF FT (BFT) interacts with FD in the nucleus, interfering with FT function under high salinity and inhibiting *AP1* expression, thereby delaying flowering (Ryu et al., 2014). TFL1 differs from FT only in 39 non-conserved amino acids but as mentioned above has an opposite biological function: TFL1 represses flowering while FT is a floral promoter (Ahn et al., 2006). It has been demonstrated that substitutions of a single amino acid (TFL1-H88; FT-Y85) or exchange of the segment B encoded by the fourth exon are sufficient to impose TFL1-like activity onto FT, and *vice versa* (Ahn et al., 2006; Hanzawa et al., 2005; Ho and Weigel, 2014). Similar to FT, TFL1 also interacts with FD, both in yeast-2-hybrid assays as well as in plant nuclei (Hanano and Goto, 2011; Wigge et al., 2005). Together, these findings suggest that activating FD-FT and repressive FD-TFL1 complexes compete for binding to the same target genes (Ahn et al., 2006). Support for this hypothesis stems from the observation that TFL1 apparently acts to repress transcription (Hanano and Goto, 2011) whereas FT seems to function as a transcriptional (co-) activator (Wigge et al., 2005). However, evidence that these protein complexes in fact share interactors such as 14-3-3 proteins or control the same targets remains sparse.

FD has been reported as direct and indirect regulator of important flowering time and floral homeotic genes such as *SUPPRESSOR OF OVEREXPRESSION OF CONSTANS 1* (*SOC1*), *SQUAMOSA PROMOTER BINDING PROTEIN-LIKE 3* (*SPL3*), *SPL4*,*SPL5*, *LEAFY* (*LFY*), *AP1*, and *FRUITFULL* (*FUL*). Several flowering time pathways contribute to *SOC1* regulation. Indeed, it has been proposed that expression of *SOC1* can be directly promoted by the FD-FT complex (Lee and Lee, 2010). However, *SOC1* expression can also be activated independently from FD-FT probably through the SPL3, SPL4, and SPL5 proteins (Lee and Lee, 2010; Moon et al., 2003; Wang et al., 2009), which have been shown to be directly or indirectly activated by the FD-FT complex (Jung et al., 2012). The activation of floral homeotic genes such as *AP1* and *FUL* in response to FD-FT activity at the SAM can at least in part be explained by the direct activation of the floral meristem identify gene *LFY* through SOC1 (Jung et al., 2012; Moon et al., 2005; Yoo et al., 2005). In addition, it has also been proposed that FD-FT complex can promote the expression of *AP1* and *FUL* by directly binding to their promoters (Abe et al., 2005; Teper-Bamnolker and Samach, 2005; Wigge et al., 2005). Taken together, these results support a central role for FD in integrating different pathways to ensure the correct timing of flowering. However, FD targets have not yet been identified at the genome scale, nor has the requirement for protein complex formation for FD function in *A. thaliana* been systematically addressed.

Here we identify direct and indirect targets of FD at the genome scale using ChIP-seq and RNA-seq in wildtype as well as in *ft-10 tsf-1* double mutants. This demonstrates that FD can bind to DNA *in vivo* even in the absence of FT/TSF. However, FD binding to a subset of targets, which includes many important flowering time and floral homeotic genes, was reduced in the *ft-10 tsf-1* double mutant, strongly supporting a role for FT/TSF in modulating FD DNA binding and expression of functionally important target genes. In addition, we report the effects of FD phosphorylation on protein complex formation with FT and TFL1 via 14-3-3 proteins *in vitro* and show how phosphorylation of FD affects flowering time *in planta*. Finally, our ChIP-seq experiments identified hundreds of previously unknown FD target genes, both in the PCCs as well as at the SAM. For example, we observed that FD directly binds to and regulates genes in hormone signaling pathways. These newly identified FD target genes represent a precious resource not only to enhance our knowledge of the photoperiod pathway but also to better understand the integration of different signaling pathways at the transcriptional level. Taken together, our findings support a role for FD as a central integrator of flowering time and provide important novel data to guide future research on the integration of diverse signaling pathways at the SAM.

## RESULTS

### FD binds G-box motives when expressed in PCCs

FD is normally expressed at the shoot apical meristem (SAM) whereas its interaction partner FT is expressed in leaf phloem companion cells (PCC). As most 14-3-3 proteins are ubiquitously expressed at moderate to high levels and have also been detected in PCCs (Deeken R. et al., 2008; Schmid et al., 2005), we reasoned that expression of FD from the PCC-specific *SUC2* promoter would maximize FAC complex formation and enable us to investigate the role of FT in modulation of FD transcriptional activity. We performed ChIP-seq on independent biological duplicates in a stable *pSUC2::GFP:FD* reporter line in Col-0 background using *pSUC2::GFP:NLS*, in which the GFP protein is fused to the nuclear localization signal (NLS), as a control. A total of 2068 and 3236 genomic regions showing significant enrichment (peaks) were identified in the first and second replicate, respectively (Fig. S1A). Overlapping results from the two biological replicates identified 1754 high-confidence peaks shared in both experiments (Fig. S1 A, Supplemental Data Set 1) and only this subset of peaks, which include important flowering time and flower development genes such as *AP1*, *FUL*, *LFY*, *SOC1*, *SEP1*, *SEP2*, *SEP3*, was used for further analysis. In both replicates, the majority of the peaks mapped to promoter regions (65,1% and 63.8%, respectively), followed by intergenic regions (16% and 16.8%), transcriptional terminator sites (9.2% and 10.7%), exons (6.4% and 5.6%) intron (2.4% and 2.3%), 5’-UTR (0.5% and 0.3%), and 3’-UTR (0.4% and 0.5%) (Fig. 1A). The relative enrichment of peaks mapping to promoter regions is in agreement with what is expected from a transcriptional regulator. In both replicates, the majority of the peaks are located between 600 bp and 300 bp upstream the nearest transcription start site (TSS) (Fig. S1D, G). *De novo* motif analysis using MEME-ChIP (Machanick and Bailey, 2011) revealed that peak regions showed a strong enrichment of G-boxes (CACGTG), which is a canonical bZIP binding site (Fig. S1J). The subset of 1754 peak regions was associated with 1676 unique genes, with 68 genes containing more than one inferred FD binding site. Taken together, these results demonstrate that, when misexpressed in the PCCs, FD is capable of binding to G-box elements in a large number of genes that are involved in diverse aspects of the plant life cycle.

**Figure 1.**
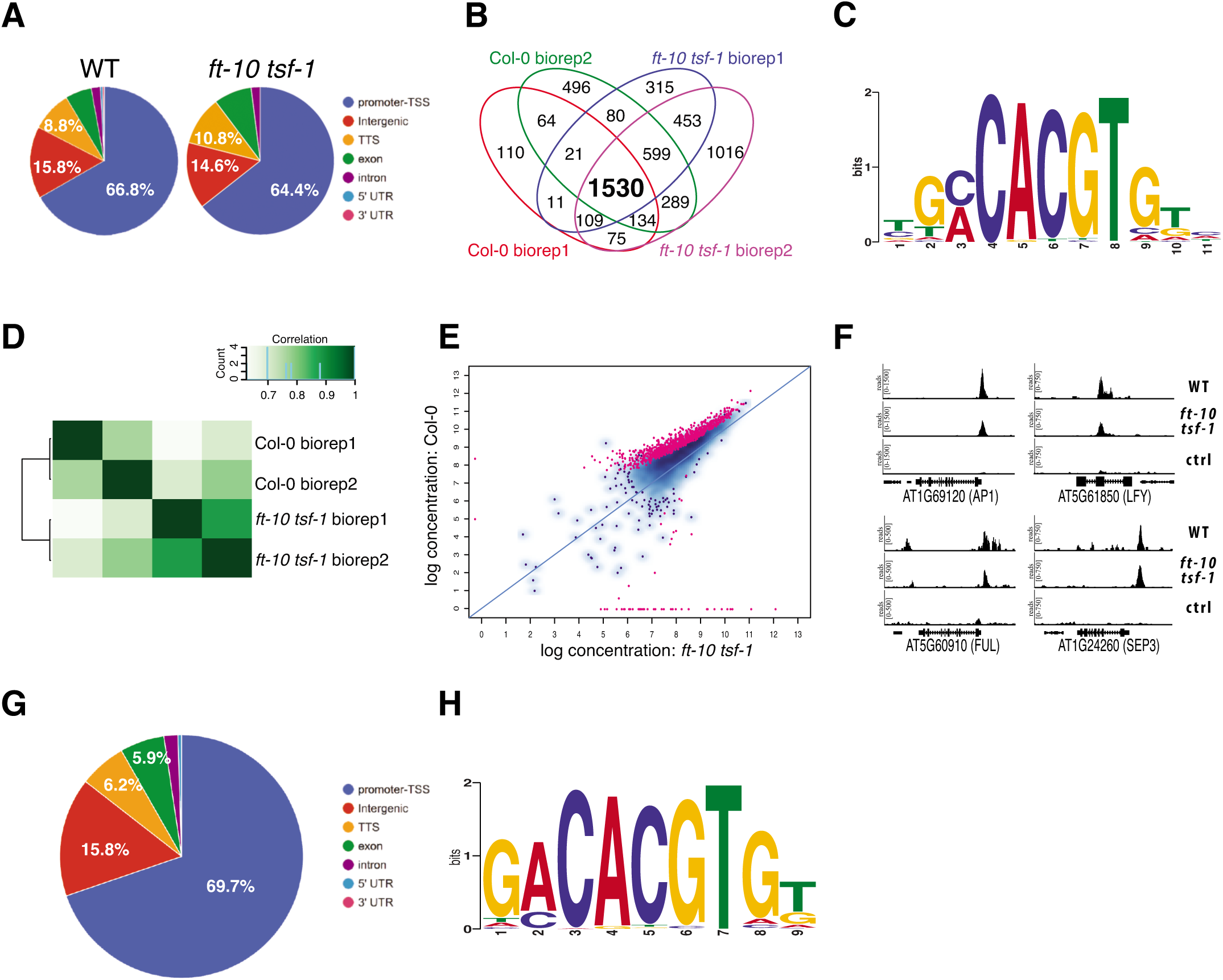
Identification of FD targets by *pSUC2::GFP:FD* ChIP-seq in WT and *ft-10 tsf-1* and *pFD::GFP:FD* ChIPseq in **(A)**Annotation of high-confidence peaks found in two biological replicates in WT and *ft-10 tsf-1*. **(B)**4-set venn diagram representing the overlapping peaks among all the biological replicates from WT and *ft-10 tsf-1*. The majority of peaks (1530) is shared between the two genetic backgrounds. **(C)**Nucleotide logo of the predicted FD binding site. **(D)**Binnding matrix (affinity scores) based on ChIP-seq reads counts for WT and *ft-10 tsf-1* samples. The presence of and TSF is sufficient to discriminate the two genetic backgrounds. **(E)**Differential bound (DB) peaks between WT and *ft-10 tsf-1*. Red dots indicate differentially bound peaks with a R < 0.05. **(F)**Reads from WT, *ft-10 tsf-1* and control sample mapped against selected flowering related genes. **(G)**Annotation of high-confidence peaks identified by ChIP-seq in two biological replicates in *pFD::GFP:FD fd-2*. **(H)**Nucleotide logo of the predicted FD binding site at the SAM.

### FT and TSF enhance binding of FD to DNA

To test whether FT and its paralog TSF are required for FD to bind to DNA, the *pSUC2::GFP:FD* reporter and *pSUC2::GFP:NLS* control constructs were transformed into the *ft-10 tsf-1* mutant background. Results from two independent biological replicates show that FD is capable of binding to DNA even in the absence of *FT* and *TSF*. Most peaks (63% and 62.1% in the first and second biological replicate, respectively) mapped to promoter regions within 600 bp and 300 bp nucleotides upstream the nearest TSS (Fig. S1B, E, H). Overall, these results are very similar to those observed for *pSUC2::GFP:FD* in Col-0 (Fig. 1A, Fig. S1B, E, H, K). Comparison between the two biological replicates identified 2696 common peaks in *ft-10 tsf-1* mutant that mapped to 2504 unique genes (Fig. S1B, Supplemental Data Set 2). Surprisingly, overlapping the sets of genomic regions bound by FD with high-confidence in WT (1754) and *ft-10 tsf-1* (2696) identified 1530 shared peaks (Fig. 1B, Supplemental Data Set 3), suggesting that FD is capable of binding to most of its targets in the absence of FT and TSF. Analysis of the sequence under the 1530 shared peaks revealed that FD maintained its strong preference for binding to G-box motifs (Fig. 1C).

Analysis of differential bound (DB) regions revealed that, although FT and TSF were not required for FD to bind DNA, their presence increased the strength of the binding and this was sufficient to discriminate the two genetic backgrounds (Fig. 1D). A total of 885 DB regions with a FDR < 0.05 were found between WT and *ft-10 tsf-1* and almost all of these loci showed higher enrichment in WT (Fig. 1E, Supplemental Data Set 4). Interestingly, this subset includes important floral homeotic genes such as *AP1*, *SEP1*, *SEP2*, and *FUL*, as well as two members of the *SPL* gene family, *SPL7* and *SPL8*. We also found FD bound to the second exon of *LFY*, a master regulator of flower development (Fig. 1F). In addition, we detected binding to loci encoding genes involved in the regulation of gibberellic acid biosynthesis and degradation such as *GA2OX4*, *GA2OX6*, and *GA3OX1* as well as to three key components of the circadian clock, *CCA1*, *LHY*, and *TIC* (Supplemental Data Set 4).

To test the robustness of our results and any possible bias due to the different genetic backgrounds used as controls, Col-0 and *ft-10 tsf-1*, peaks were called again using *pSUC2::GFP:NLS* in Col-0 as single negative control. Analysis identified 917 DB (Fig. S2), which is comparable to the 885 DB genes from the previous analysis (Fig. 1E). In addition, affinity test analysis clustered by genotype rather than the control used (Fig. S2), ruling out a bias due to the usage of different genetic backgrounds for peak calling. Importantly, FD is capable of inducing the known FAC target gene *AP1* in leaves when expressed under the *pSUC2* promoter, suggesting that a functional FAC can be formed in the phloem companion cells when FD is present (Fig. S3A). The finding that *AP1* expression could only be observed in the Col-0 background but not in *pSUC2::GFP:FD ft-10 tsf-1* further supports this interpretation. However, in contrast to *AP1*, we failed to detect induction of *SOC1* in the PCCs of *pSUC2::GFP:FD* (Fig.S3A), suggesting that other co-factor(s) that are probably specifically expressed at the SAM might be required to fully activate FD target gene expression.

### FD phosphorylation is required for complex formation and to promote flowering

To verify the binding of FD to G-boxes *in vitro* we performed electrophoretic mobility shift assays (EMSA) using the bZIP domain of the *A. thaliana* FD protein (FD-C) and a 30bp fragment from the *SEP3* promoter containing a G-box that we had identified as FD target region in our ChIP-seq (Fig. 1F) as a probe. We observed weak binding of FD-C, but failed to detect higher order complexes when 14-3-3, FT, or both were added (Fig. 2A). In contrast, a clear supershift with 14-3-3 and FT was observed when a phosphomimic variant of FD-C, FD-C_T282E, was used (Fig. 2B). Interestingly, TFL1, which is similar to FT in structure (Ahn et al., 2006) but delays flowering, was capable of forming a complex with 14-3-3 and wildtype FD-C (Fig. 2A). Similar results were obtained with the full-length version of FD (Fig. S4A). Taken together, these results demonstrate that *A. thaliana* FD is capable of binding to DNA without FT, confirming results from our ChIP-seq experiments. Furthermore, our results suggest that the unphosphorylated form of FD, in complex with 14-3-3 proteins, can interact with TFL1.

**Figure 2.**
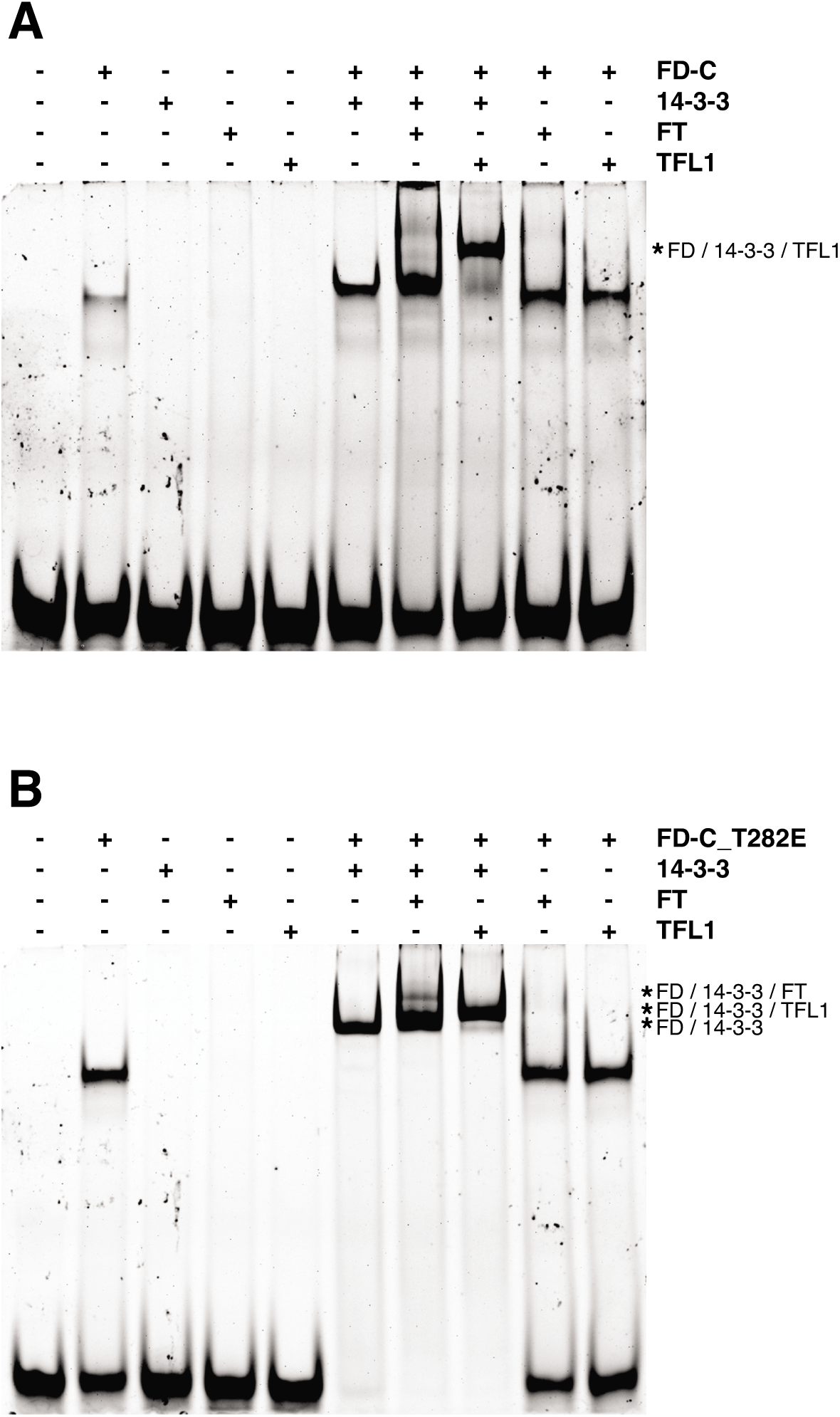
The C-terminal part of FD (FD-C) binds to a G-box in the SEP3 promoter *in vitro*. **(A)**Electrophoretic mobility shift assay (EMSA) of the wild-type form of FD-C in combination with 14-3-3v, FT, and TFL1. FD-C weakly binds the probe on its own but it is not able to form complex with 14-3-3v and FT. However, FD-C forms a complex with 14-3-3? and TFL1 capable of binding the G-box. **(B)**Phosphomimic version of FD-C (FD-C_T282E) in combinations with 14-3-3v, FT and TFL1. The phosphomimic version of FD-C binds the G-box alone and it is interacting with 14-3-3v, which facilitates interaction with FT and TFL1. Both, wild-type and phosphomimic version of FD-C, require 14-3-3v for interaction with FT or TFL1. Asterisk (*) indicate shifted probe.

To investigate the importance of FD phosphorylation *in vivo* we complemented the *fd-2* mutant with *pFD::FD*, *pFD::FD-T282E*, and *pFD::FD-T282A* (which cannot be phosphorylated) and determined flowering time of homozygous transgenic plants. Plants transformed with the WT version of FD rescued the late flowering phenotype of *fd-2*, indicating that the rescue construct was fully functional. In contrast, plants transformed with the T282A version flowered with the same number of leaves as *fd-2*, demonstrating that FD needs to be phosphorylated to induce flowering. Interestingly, plants transformed with the T282E phosphomimic version of FD flowered even earlier than WT (Fig. 3), indicating that control of FD phosphorylation is important for its function *in vivo*. To test whether serine 281 (S281), which is located next to T282, constitutes a potential FD phosphorylation site, we complemented *fd-2* with *pFD::FD-S281E* and *pFD::FD-S281E/T282E* constructs. Interestingly, these lines flowered as early as plants transformed with the phosphomimic version T282E (Fig. 3), indicating that S281 could be a possible FD phosphorylation site but that double-phosphorylation of S281/T282 does not accelerate flowering any further. These *in vivo* results are in agreement with our EMSA results and confirm that phosphorylation of FD is required for its function and needs to be finely regulated in order to avoid either premature or delayed flowering. It should be noted, however, that the phosphomimic version of the C-terminal fragment of FD (as used in the EMSA analyses) is insufficient to fully rescue the late flowering of *fd-2* (Fig. S3B), suggesting that the N-terminal region of FD, even though it does not contain any known functional domains, nevertheless contributes to FD function.

**Figure 3.**
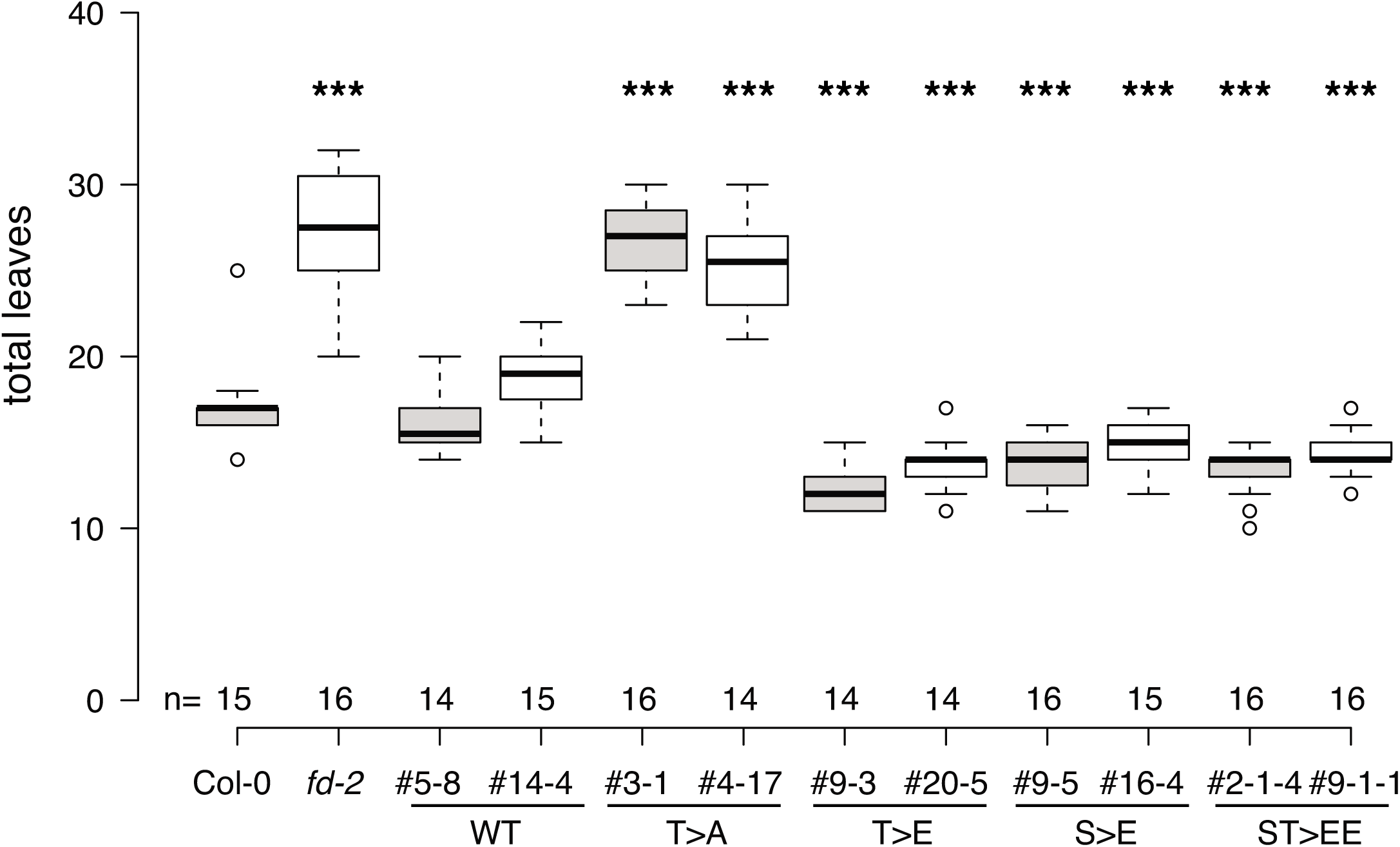
Phosphorylation of FD at threonine 282 T282) modulates flowering time in *A. thaliana*. Expression of wildtype (WT) *pFD::FD* rescues the late flowering phenotype of *fd-2*. Mutation of T282 to alanine (T>A) in *pFD::FD_T282A*, which prevents phosphorylation, abolishes rescue of *fd-2*. Mutations mimicking constitutive phosphorylation of T282 (T>E), S281 (S>E), or both (ST>EE) induce early flowering. Results are shown for two independent homozygous lines per construct. Statistical significance was calculated using unpaired t-test compared to Col-0. *** indicate a significance level p < 0.01.

### Targets of FD at the SAM

The rationale for carrying out the initial ChIP-seq experiments in PCCs was to maximize the likelihood of FAC formation and to study the contribution of FT/TSF to FD DNA binding. However, since our ChIP-seq and EMSA results indicated that FD-FT interaction is not required for FD to bind to DNA, we decided to determine direct targets of FD in its natural context at the SAM.

To this end we performed ChIP-seq using a *fd-2* mutant that had been complemented using a *pFD::GFP:FD* construct (Fig. S3C). ChIP-seq was performed using two independent biological replicates from apices of 16-day-old plants grown in LD condition. In the two replicates, we could identify 703 and 1222 FD-bound regions, respectively, of which 595 were shared between the replicates (Fig. S1C, Supplemental Data Set 5). Of these, 69.7% mapped to core promoter regions within 300 to 600 bp upstream of the nearest TSS, 15.8% in intergenic regions, followed by TTS (6.2%), exons (5.9%), introns (1.8%) and 5’-UTRs (0.5%) (Fig. 1G, Fig. S1F, I). Similar to the situation in our PCC-specific ChIP-seq analyses we found a G-box as the most overrepresented transcription factor binding site under the peak region (Fig. 1H, Fig. S1L). The 595 peak regions shared between the replicates mapped to 572 individual genes, which we consider high-confidence *in vivo* targets of FD at the SAM and which include important flowering-related genes such as *AP1*, *FUL*, *SOC1*, and *SEP3*. The precise location of the FD binding site in the *AP1* promoter has been discussed controversially (Benlloch et al., 2011; Wigge et al., 2005). Taking into account all six ChIP-seq datasets, we were able to extract a 64 bp sequence covering the peak summits on the *AP1* promoter (Fig. 4A,B). Interestingly, this sequence lies about 100 bp downstream of a C-box that had previously been implicated in FD binding to *AP1* (Wigge et al., 2005), but contains several palindromic sequences. However, none of them is a *bona fide* G-box. We selected three potential binding sites within the 64 bp sequence and tested them, along with the upstream C-box, by EMSA for FD binding (Fig. 4C, S4B). Results show that only the phosphomimic version of FD-C (FD-C_T282E) in combination with 14-3-3 can bind to DNA. Furthermore, a supershift is detected for all palindromic sites tested, included the C-box, when TFL1 is added. In contrast, for FT an additional shift resembling the pattern obtained with the G-box in *SEP3* promoter was only observed for “site 2” (Fig. 4C, 2B). Closer inspection of the nucleotide sequences of the probes used for the G-box in the *SEP3* promoter and the “site 2” in the *AP1* promoter revealed that the possible FD binding site in the *AP1* promoter (GTCGAC) is also present in the *SEP3* promoter, where it overlaps with the G-box (Fig. 4D). Interestingly, in the context of the *SEP3* probe, full-length FD and FD-C tolerated mutating the core of the G-box from CG to GC, whereas CG to TA mutations as well as converting the G-box to a C-box (GACGTC) abolished binding *in vitro* (Fig. S4D). To further test the site 2 on *AP1* promoter as real binding site of FD, we mutated its core from CG to TA and checked whether this was sufficient to abolish the FD binding. Results show that indeed the binding of FD was strongly abolished except in the presence of TFL1 (Fig. S4E).

**Figure 4.**
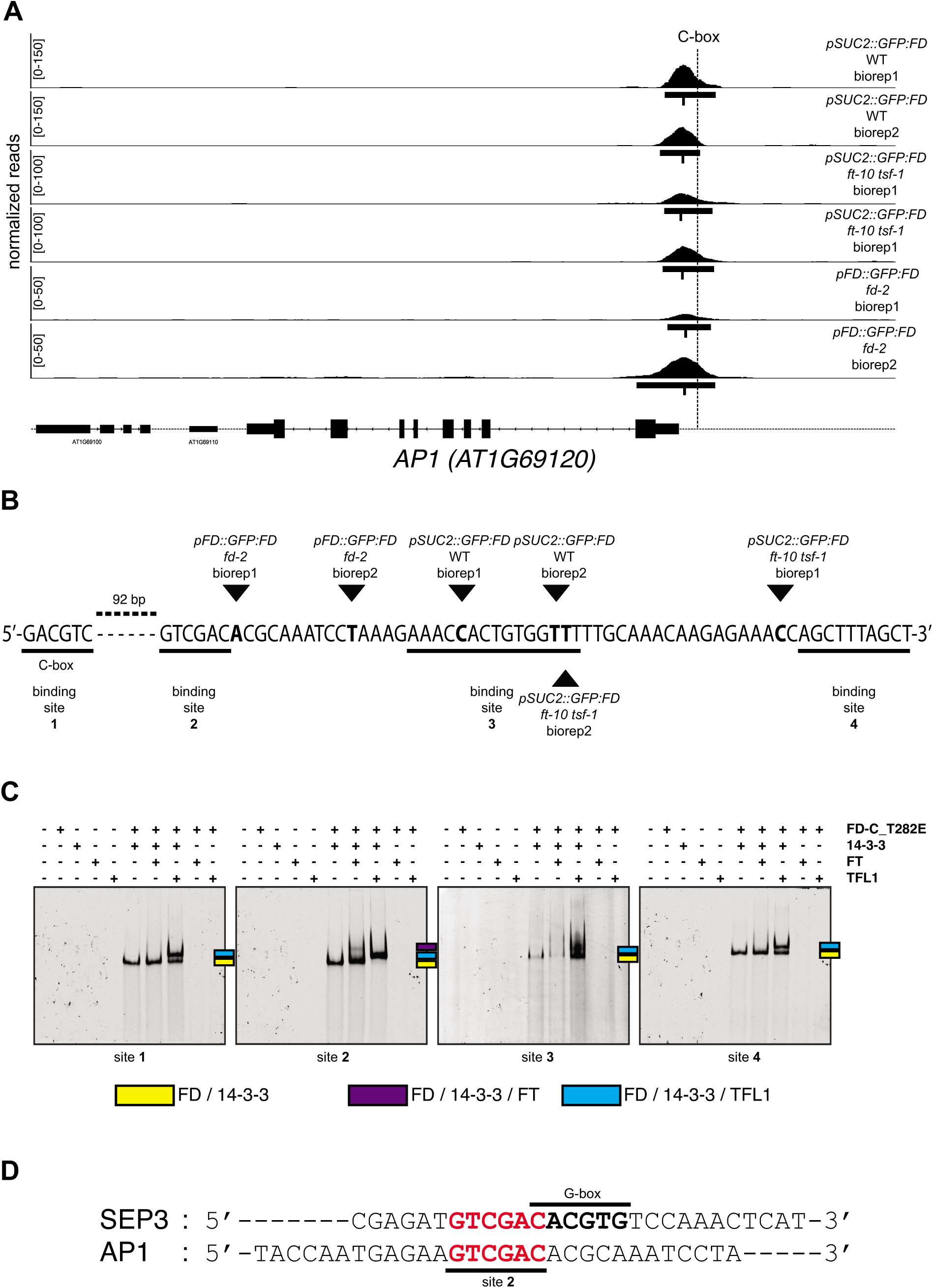
Mapping of the FD binding site in the AP1 promoter. **(A)**Normalized reads from six ChIP-seq experiments mapped on the *AP1* locus. The result shows that the C-box is laying upstream of all peak summits. **(B)**Nucleotide sequence encompassing the six peak summits shows several palindromic regions representing putative binding sites of FD on *AP1* promoter. The distance between the closest potential FD binding site under the ChIP-seq peaks and the C-box is 92 bp. Black triangles indicate the summits of the six separate ChIP-seq experiments. Putative FD binding sites are underlined and numbered from 1 to 4. **(C)**Electrophoretic mobility shift essay (EMSA) of the phosphomimic version of FD-C (FD-C_T282E) in combinations with 14-3-3v, FT and TFL1 using the four putative binding sites reported in panel B. Free probes are not visible because gels were running longer to maximize the distance between shifted probes. Coloured squares indicate shifted probes. **(D)**Comparison of the probes used for EMSA: the G-box in *SEP3* promoter (Fig. 2) and the binding site 2 in *AP1* promoter. The putative FD binding site in *AP1* promoter is also conserved in *SEP3* promoter and it is overlapping with the G-box.

Take together our findings exclude the C-box as the FD binding site in the *AP1* promoter. Furthermore, our results suggest that FD can bind other motifs as well, possible through interaction with interaction partners other than 14-3-3 and FT/TSF, and we characterized a new binding site (GTCGAC) that could be the most likely real FD binding site in *AP1* promoter.

### Differentially expressed genes at the SAM and direct targets of FD

To test which of the 595 high confidence targets we had identified by ChIP-seq at the SAM were actually transcriptionally regulated by FD we performed RNA-seq on apices from *fd-2* mutant and the *pFD::GFP:FD fd-2* rescue line. 21 day-old SD-grown seedlings were shifted to LD to synchronize flowering and apices were harvested on the day of the transfer to LD (T0), as well as 1, 2, 3, and 5 days after the shift (T1, T2, T3, T5) from three independent biological replicates.

Differentially expressed (DE) genes were called for each time point and genes with an adjusted p-value (padj) lower than 0.1 were selected as significantly DE. In total 1759, 583, 2421, 924, and 153 DE genes were identified in T0, T1, T2, T3, and T5, respectively, corresponding to 4189 unique genes (Fig. 5A, Supplemental Data Set 6). PCA analysis showed that the first and second principal component, which explain 37% and 21% of the total variance, corresponded to the different time points and genotypes, respectively (Fig. S5A). The best separation between the genotypes in the PCA was observed at T3 and T5, indicating that FD contributes to the transcriptional changes at the SAM mainly after exposure to two long days. This observation is in agreement with the expression profile of FD, which in the *pFD::GFP:FD* rescue line increased after T2 (Fig. S5B). In contrast, FD expression remained low in the *fd-2* mutant, indicating the validity of our experimental approach (Fig. S5B).

**Figure 5.**
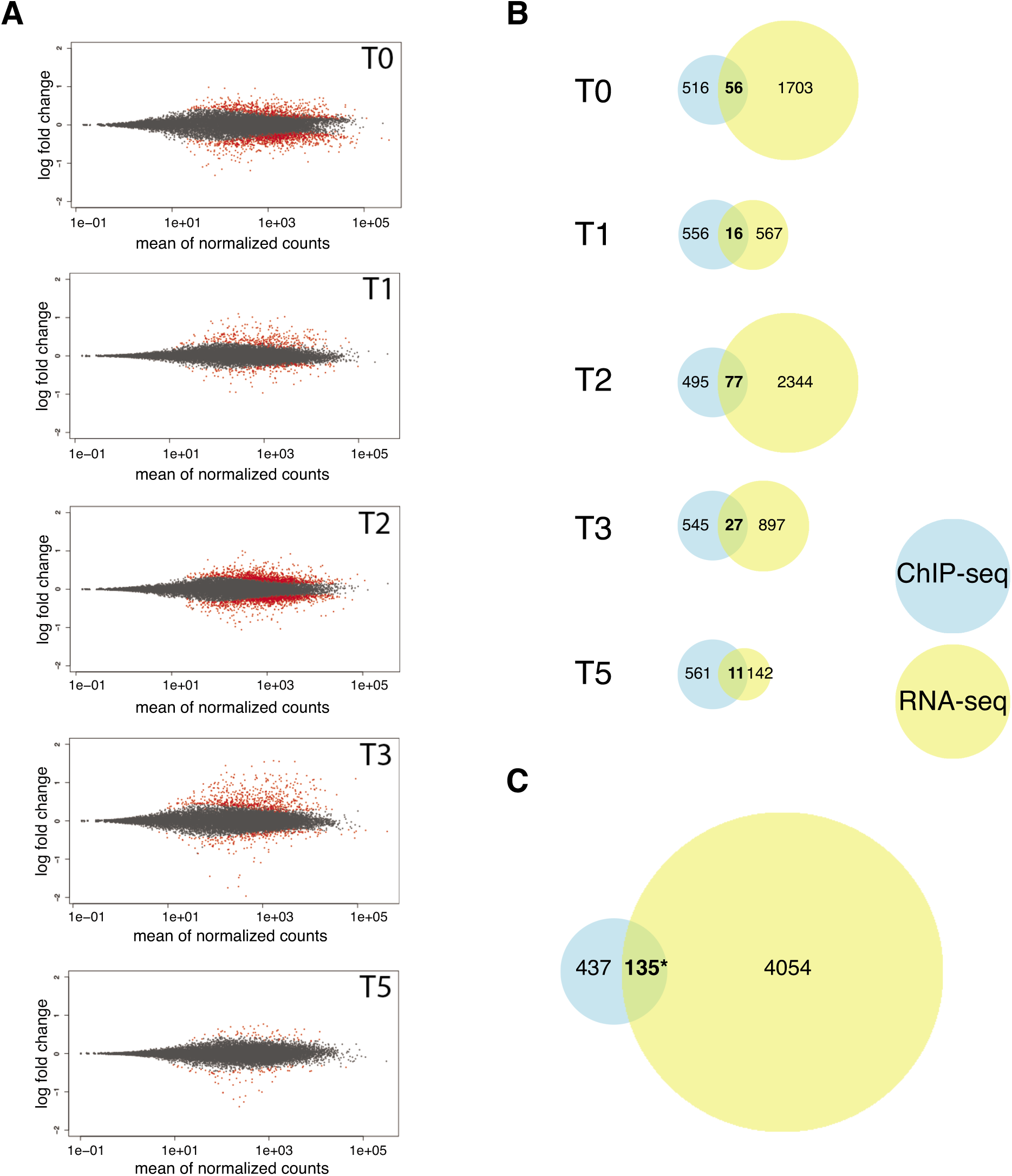
RNA-seq results at the shoot apical meristem. **(A)**Scatter blot of differential expressed (DE) genes between the *fd-2* mutant and *pFD::GFP-FD fd-2* (control) at 5 ime points before and during the transition to photoperiod-induced flowering. T0 – T5 indicate day of sample collection before (T0) and 1, 2, 3, 5 days after shifting plants to long day. Red dots indicate DE genes with a padj < 0.1. **(B)**Venn diagrams showing the overlap between FD target genes identified by ChIP-seq and DE genes found by RNA-seq at the SAM at each time point. **(C)**enn diagram showing the overlap between FD target unique genes identified by ChIP-seq and DE unique genes in at least one time point found by RNA-seq at the SAM. A total of 135 genes were classified as putative direct targets of FD. Statiscal significance was calculated using the Fisher’s exact test. Asterisk (*) indicates a significance level p = 1.03E-07.

Next, we intersected the list of genes that were bound by FD at the SAM (572) with the list of DE genes (4189). In total, 135 (23.6%) of the 572 FD-bound genes were significantly DE at the SAM during the transition to flowering at least at one timepoint, indicating that these genes are transcriptionally regulated by FD, which is more than expected by chance (Fig. 5B, C, Supplemental Data Set 7). Among the 135 directly bound and differentially expressed FD targets we observed several previously known FD-regulated flowering time and floral homeotic genes including *AP1*, *FUL*, and *SOC1* (Fig. 1F, S6A). In addition, this set of high-confidence FD targets contained also the MADS box gene *SEP3*, the promoter of which is bound by FD and which is down-regulated in *fd-2* mutant (Fig. 1F, S6A). Interestingly, we did not observe binding of FD to any of the other members of *SEPALLATA* gene family in ChIP-seq samples from the SAM, although we did detect FD binding in promoter regions of *SEP1* and *SEP2*, but not *SEP4*, in ChIP-seq from seedlings in which FD had been misexpressed from the *SUC2* promoter. One possible explanation for this is that the ChIP-seq at the SAM apparently worked less efficiently and identified fewer FD targets (1754 vs. 595), which might result in a larger number of false negatives. In agreement with this interpretation, *SEP1* is down-regulated in *fd-2* mutant (Fig. S6), indicating that FD directly or indirectly regulates the expression of *SEP1* at the SAM. Interestingly, we also found FD bound to *TPR2*, a member of the *TOPLESS (TPL)-related* gene family. TPL and its family members (TPR1, TPR2, TPR3 and TPR4) are strong transcriptional co-repressors and they interact with other proteins throughout the plant to modulate gene expression (Causier et al., 2012). *TPR2* is down-regulated in the *fd-2* mutant throughout floral transition from T0 to T5 (Fig. S6), indicating FD might regulate development at the SAM through *TPR2* in a photoperiod-independent manner. Gene Ontology (GO) analysis of these 135 genes that were bound and differentially expressed by FD revealed significant enrichment in several biological process categories (Fig. S7), including “flower development” and “maintenance of inflorescence meristem identity”, as one might expected for a flowering time regulator such as FD. More surprisingly, however, genes related to the “response to hormone” category were also significantly overrepresented (Supplemental Data Set 8). Among these 27 genes are four genes best known for their role in jasmonate signaling (*MYC2*, *JAZ3*, *JAZ6* and *JAZ9*), three genes directly connected to auxin signaling (*ARF18*, *WES1*, and *DFL1*), four genes involved in abscisic acid signaling (*ALDH33I1*, *ATGRDP1*, *HAI1* and *PP2CA*), and the flowering-related gene *SOC1*, which is well-known to be regulated by gibberellins (Supplemental Data Set 8). Closer inspection of the expression profiles of these 27 candidate genes revealed that *ARF18* showed a trend similar to *SOC1*, being strongly induced after T2 in Col-0 but not in *fd-2*. The four jasmonate-related genes showed a peculiar expression profile in *fd-2*, *i.e.* an increase from T0 to T1, decrease in T2, another increase in T3, and decreasing in T5. Since this peculiar expression profile was observed in three *JAZ* genes, we checked the remaining genes in this family and found that 11 out of 13 displayed the same pattern (Fig. S6). Furthermore, this profile was also observed in three other genes (*DMR6*, *ESP* and *TOE2*), all of which have previously been implicated in pathogen resistance and the jasmonate pathway (Fig. S7B). Taken together, these results suggest that FD plays an active role not only in the regulation of flowering time but also functions as a hub for different hormone signaling pathways.

### Validation of FD targets

We selected a subset of putative FD direct target genes and determined their expression in early flowering FD overexpression lines (*p35S::FD*) and Col-0. To minimize any bias due to the early flowering of *p35S::FD*, experiments were carried out in vegetative 7-day-old LD-grown seedlings. For validation, we selected genes known to play a major role in floral transition, genes that according to Gene Ontology are involved in flowering time and floral development, and other genes that showed a marked differential expression in *fd-2* but for which a role in flowering time regulation had not previously been studied in detail. qRT-PCR assays confirmed that both *SOC1* and *AP1* were strongly up-regulated in *p35S::FD* (Fig. 6). Although we had only found *SEP3* to be bound by FD in the SAM ChIP-seq analysis, we tested expression of all four *SEPALLATA* genes (*SEP1 – SEP4*) in the *p35S::FD* line. *SEP3* was the only *SEP* gene that was strongly induced in seedlings in response to *FD* overexpression, while *SEP1* and *SEP2* showed only moderate induction. In contrast, expression of SEP4 did not show difference between *p35S::FD* and Col-0 (Fig. 6). Interestingly, *SEP1*, *SEP2*, and *SEP3* were also bound by FD in PCC-specific ChIP-seq in seedlings and *SEP1* and *SEP3* displayed strong DE in RNA-seq (Fig. S6). *AS1*, which has been demonstrated to be involved in flowering time by regulation of *FT* expression in leaves (Song et al., 2012), did not show significant difference in expression between Col-0 and *p35S::FD*. We also tested two FRIGIDA-like genes, *FRI-like 4a* and *FRI-like 4b*, of which *FRI-like 4b* showed a decreased expression in *p35S::FD*. In addition, we also tested two genes, *MYC2* and *AFR1*, which were bound by FD in both the *pSUC2* and *pFD* ChIP-seq experiments, differentially expressed at the SAM, but not differentially bound in *ft-10 tsf-1* mutant, *i.e.* not directly influenced by the presence of FT and TSF, for their contribution to flowering time regulation. *MYC2* showed no differences in expression in *p35S::FD* compared to Col-0, whereas *AFR1* was up-regulated in *p35S::FD* (Fig. 6). To genetically test the role of these two genes in the regulation of flowering we isolated T-DNA insertion lines and determined their flowering time under LD at 23°C. Both *myc2* and *afr1* were significantly early flowering, both as days to flowering and total leaf number, compared to WT (Fig. 7), confirming their role in regulating the floral transition.

**Figure 6.**
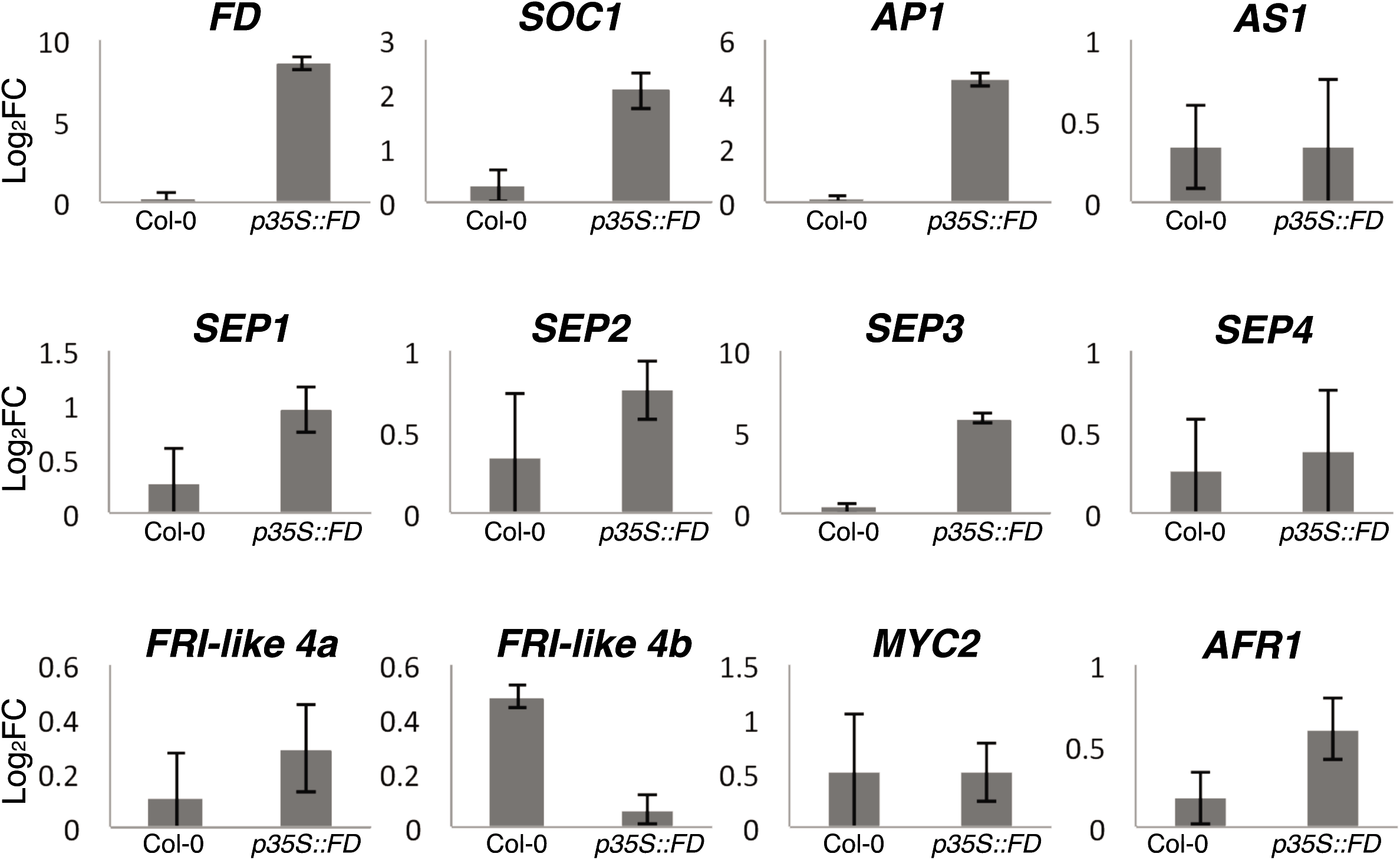
Valilidation of FD targets in Col-0 and *p35S::FD*. qRT-PCR analysis of 12 putative direct targets of FD. RNA was isolate from 7 days old seedlings to minimize any o the early flowering of the *p35S::FD* line. Error bars represent ±SD from three biological replicates.

**Figure 7.**
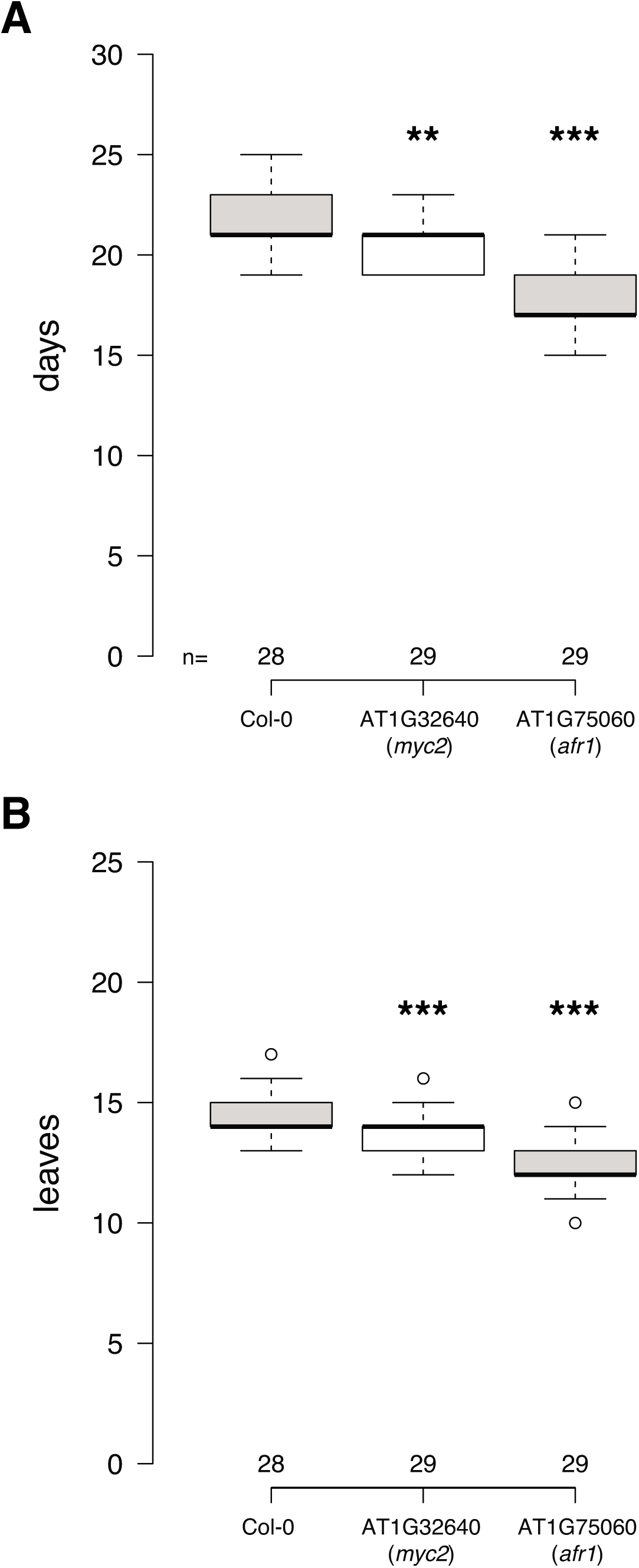
Flowering time of *myc2* and *afr1*. Flowering time of homozygous of *myc2* and *afr1* T-DNA insertion lines was scored as days to flowering **(A)** and total leaves **(B)**. Statistical significance was calculated using unpaired t-test compared to Col-0. *** and ** indicate a significance level p < 0.01 and p < 0.05, respectively.

## DISCUSSION

*FD* was originally identified as a component of the photoperiod-dependent flowering pathway in *A. thaliana* based on the late flowering phenotype of the loss-off-function mutant (Koornneef et al., 1991). *FD*, which encodes a bZIP transcription factor, is expressed in the SAM prior to floral transition but does not induce flowering alone. Later, it was demonstrated that FD physically interacts with FT, the florigen, and that this interaction is important for its function as a promoter of flowering (Abe et al., 2005; Wigge et al., 2005). In addition, FD was found to also interact with TFL1, which is normally expressed in the SAM and antagonizes the function of FT as floral activator. This and other findings led to the hypothesis that FD is held in an inactive state through TFL1 interaction in the vegetative SAM. When FT is induced in the PCCs and transported to the SAM in response to inductive photoperiod, FT competes with TFL1 for interaction with FD, eventually resulting in the formation of transcriptionally active FD-FT complexes (Ahn et al., 2006). However, the exact molecular mechanisms of FD action and its genome-wide targets remained largely unknown. Here we employed biochemical, genomic, and transcriptomic approaches to clarify the role of FD in the regulation of flowering transition in *A. thaliana*.

We found that neither FT nor TSF are required for FD to bind to DNA but that their presence increases the strength of FD binding on a subset of target loci, which encode known flowering time and floral homeotic genes such as *AP1*, *SEP1*, *SEP2*, and *FUL*. Our data are compatible with the model described by (Ahn et al., 2006), according to which FT acts as a transcriptional coactivator. Without FT, FD is still capable of binding DNA but does not seem to activate transcription. In this context, our EMSA results are of particular interest as they demonstrate that, at least *in vitro*, TFL1 is capable of interacting with unphosphorylated FD via 14-3-3 proteins, suggesting that the transcriptionally inactive ternary FD/14-3-3/TFL1 complex is the ground state at the SAM. Only after FD has been phosphorylated can FT, together with 14-3-3 proteins, form an active FAC to induce flowering. This requirement for phosphorylation of T282 of FD adds another safeguard to the system that might help to prevent disastrous premature induction of flowering. Our results clearly suggest that phosphorylation is important for FD function and add to our understanding concerning the role of FD phosphorylation, which had mostly been based on the analyses of a FD/14-3-3/Hd3a complex in rice using a short FD peptide (Kaneko-Suzuki et al., 2018; Taoka et al., 2011).

Which kinases regulate phosphorylation of FD *in vivo* has been a matter of debate, but recently two calcium-dependent kinases, CPK6 and CPK33, have been shown to phosphorylate FD (Kawamoto et al., 2015). Building on this, we show that expression of a non-phosphorable version of the FD protein (T282A) under the control of the *pFD* promoter failed to rescue the late flowering of *fd-2*. In contrast, expression of a phosphomimic version of FD (T282E) resulted in early flowering when expressed in *fd-2*. Similar results were obtained using a S281E phosphomic FD. These results indicate that the phosphorylation of FD must be tightly controlled to prevent premature flowering. Interestingly, both CPK6 and CPK33 are more strongly expressed in transition apices than they are in vegetative apices (Schmid et al., 2005), which would be in agreement with an activation of FD by these two kinases during floral induction. Somewhat surprisingly we observed that the C-terminal part of the FD protein, which includes the bZIP domain and the phosphorylation site, was sufficient to trigger complex formation with FT (and TFL1) and 14-3-3 proteins. This suggests that the N-terminal region of FD, which is predicted to be highly unstructured and contains a stretch of 25 amino acids containing 19 serine residues, might be dispensable for FD/14-3-3/FT complex formation. However, the N-terminal region of FD is evolutionarily conserved, indicating that it may contribute to FD function. This notion is supported by our observation that expression of the C-terminal part of FD in plants only partially restored the late flowering of *fd-2* mutants.

Part of the flowering promoting activity of FD can probably be expressed through its effect on members of the *SEP* gene family of MADS-domain transcription factors, which are required for the activity of the A-, B-, C-, and D-class floral homeotic genes (reviewed in Theissen et al., 2016). In addition to its function as a floral homeotic gene, *SEP3* has also been reported to promote flowering by accumulation in leaves under FT regulation (Teper-Bamnolker and Samach, 2005) and as downstream target of the miR156-SPL3-FT module in response to ambient temperature (Hwan Lee et al., 2012). However, how *SEP3* is regulated at the SAM has remained unclear. Interestingly, we found that FD bound strongly to the *SEP3* promoter and *SEP3* is downregulated in the *fd-2* mutant. As FD also binds to the promoter and activated expression of the A-class gene *AP1*, FD activity might be sufficient to induce formation of sepals, which form the outmost floral whorl, and which according to the quartet model require the formation of a SEP/AP1 complex (Theissen et al., 2016). However, it should be noted that *fd* mutants do not display notable homeotic defects, indicating that FD is clearly not the only factor regulating *SEP3* and *AP1* expression. Furthermore, binding of FD to *AP1* is unlikely to be mediated by a C-box as previously suggested (Taoka et al., 2011; Wigge et al., 2005) as the summits of the ChIP-seq peaks do not cover this region of the *AP1* promoter. Interestingly, this region contains several palindromic sequences, one or more of which most likely mediate FD binding to the *AP1* promoter. Another interesting outcome of our analyses is that FD might contribute to the regulation of other processes in the plant besides flowering. In particular, we found that FD directly regulated the expression of genes involved in several hormone signaling pathways. For example, we observed FD binding to the promoter of *MYC2*, a bHLH transcription factor that plays a key role in jasmonate response. It has been shown that MYC2 forms a complex with JAZ proteins and the TPL co-repressor, and that this interaction is dependent on NINJA proteins (Pauwels et al., 2010). In this context it is noteworthy that FD also bound directly to the promoter of *TPR2* promoter and that *TPR2* was strongly downregulated in *fd-2*. This finding indicates that FD not only regulates MYC2 but also at least some of the interacting TPL-like transcriptional co-repressors. Finally, we also observed strong binding of FD to (and misexpression of) a number of JAZ genes in either PCCs and/or the SAM in our ChIP-seq and RNA-seq data. Taken together, this indicates that FD may control the expression of three core components of jasmonate signaling: *MYC2*, *TPR2*, and several *JAZ* genes. These results support earlier findings that had reported a link between jasmonate signaling components and flowering time regulation. JAZ proteins have been shown to regulate flowering in leaves through the direct interaction with the floral repressors TOE1 and TOE2, which is also bound by FD and differentially expressed in *fd-2*, and the regulation of FLC that negatively regulate *FT* expression (Zhai et al., 2015). Moreover, MYC2 has also been reported to affect flowering time by regulating *FT* expression in leaves (Wang et al., 2017; Zhai et al., 2015). However, previous publications had reported contradictory results concerning the flowering phenotype of the *myc2* mutant, ranging from late flowering (Gangappa and Chattopadhyay, 2010) to early flowering (Wang et al., 2009) or no obvious effect (Major et al., 2017). In our conditions the *myc2* mutant showed an early flowering time compared to Col-0, which in agreement with the report from Wang and colleagues (Wang et al., 2009) (Fig. 7). We also identified *ARF18*, a member of the auxin response factors protein family, as direct target of FD. Notably, the expression of *ARF18* is strongly induced after T2 in Col-0 but not in *fd-2* and this pattern is the same of known direct FD targets, e.g.: *AP1* and *SOC1*. Moreover, *ARF18* is also induced at the SAM during floral transition (Schmid et al., 2005) providing further evidence for a possible link between FD and *ARF18*. In summary, our findings suggest a link between the photoperiodic pathway gene FD and hormone signaling pathways. Although further experiments will be necessary to better understand this connection, we hypothesize that linking hormone signaling to flowering time through FD regulation might allow plants to fine tune their flowering time response to abiotic and biotic stresses.

Apart from connecting FD with hormone signaling we characterized another target gene in more detail. *AFR1*, which encodes a putative histone deacetylase subunit, had previously been shown to negatively affect the expression of *FT* in the leaves and *afr1* mutations cause early flowering (Fig. 7)(Gu et al., 2013). Our results suggests that FD might modulate flowering through ARF1-mediated regulation of chromatin. However, such regulation would most likely not be mediated by *FT*, as *FT* is normally not expressed at the SAM.

Taken together, our results support the role of FD as a key regulator of photoperiod-induced flowering and the expression of A- and E-class floral homeotic genes in *A. thaliana*. Furthermore, FD might play an important role in coordinating the crosstalk between the photoperiod pathway and hormone signaling pathways, and provide a convergence point for diverse environmental and endogenous signaling pathways..

## METHODS

### Plant materials and growth conditions

*Arabidopsis thaliana* accession Col-0 was used as wild-type. Mutants investigated in this study are: *fd-2* (SALK_013288), *ft-10* (GABI_290E08), *tsf-1* (SALK_087522), *myc2* (SALK_017005), *arf1* (SALK_026979) (Tab. S1). Seeds were stratified for 3 days in 0.1% agar in the dark at 4°C and directly planted on soil. Plants were grown on soil under long day (16 hours of light and 8 hours of night) or under short day (8 hours of light and 16 hours of night) at 23°C, 65% relative humidity. Plants used for flowering time measurements were grown in a randomized design to reduce location effects in the growth chambers.

### DNA vectors and plant transformation

DNA vectors used in this study are listed in table S2. Coding sequences were amplified by PCR from cDNA and cloned into either pGREEN-IIS vectors for flowering time studies or pET-M11 vectors for protein expression. Final constructs were transformed by electroporation in *Agrobacterium tumefaciens* and Arabidopsis plants of accession Col-0 and *fd-2* were transformed by the floral dip method. Basta treatment (0.1% v/v) was used for screening for transgenic lines.

### ChIP and ChIP-seq

Approximately 1.5 grams of seedlings (*pSUC2::GFP:FD; pSUC2::GFP:NLS*) or 300 mg of manually dissected apices (*pFD::GFP:FD; Col-0*) from 16 days old plants grown on soil under long day 23°C were harvested and fixed in 1% formaldehyde under vacuum for 1 hour. ChIP was performed as previously described (Kaufmann et al., 2010) with the following minor changes: sonication was performed using a Covaris E220 system (conditions: intensity 200 W, duty 20, cycles 200, time 120 seconds), incubation time with antibody was increased to over-night, incubation time with protein-A agarose beads was increased to 4 hours, purification of DNA after de-cross linking was performed with MinElute Reaction Cleanup Kit (Qiagen).

Anti-GFP from AbCam (ab290) was used for immuno-precipitation. ChIP-seq libraries were prepared using TruSeq ChIP Library Preparation Kit (Illumina) and BluePippin was used for gel size selection of fragments between 200 bp and 500 bp. Final concentration and size distribution of the libraries were tested with Qubit and BioAnalyzer (Agilent High Sensitivity DNA Kit). Libraries were sequenced on an Illumina HiSeq3000 system using the 50bp single end kit. All data are available from the accession number PRJEB24874.

### RNA extraction, RNA-seq and expression analysis

For RNA-seq, Col-0 and *fd-2* plants were grown for 21 days under short day 23°C and then shifted to long day 23°C. RNA was extracted from manually dissected apices collected the day of the shift (T0) and 1, 2, 3, 5 days after shifting (T1, T2, T3 and T5 respectively) using the RNeasy Plant Kit (Qiagen) according to manufactures instructions. RNA integrity and quantification were determined on a BioAnalyzer system. 1 µg of of RNA was used to prepare libraries using the TruSeq RNA Library Prep Kit (Illumina). All libraries were quality controlled and quantified by Qubit and Bioanalyzer and run on a Illumina HiSeq3000 with 50bp single end kit. All RNA-seq data have been deposited at the accession number PRJEB24873.

Validation of the selected FD targets was performed in 7 days old seedlings grown on soil under long day at 23°C.

RNA was extracted using the RNeasy Plant Kit (Qiagen) according to manufactures instructions. cDNA was synthetize using the RevertAid RT Reverse Transcription Kit (ThermoScientific) according to the manufacture instructions. qRT-PCRs were performed on a CFX96 Touch Real-time PCR Detection System (BioRad) using LightCycler 480 SYBR Green I Master (Roche). Oligonucleotides used as primers for qRT-PCR are listed in table S3.

### ChIP-seq and RNA-seq analysis

Raw data from ChIP-seq were trimmed of the adapters and aligned to the *A. thaliana* genome (TAIR10 release) using bwa (Li and Durbin, 2010). MACS2 was used to call peaks using default parameters (Zhang et al., 2008). Mapped reads from samples expressing GFP:NLS under the same promoter of the GFP:FD (*e.g.*: *pSUC2*) in seedlings experiments or Col-0 without any vector in apices experiments were used for normalization. Differential bound analyses were carried out using the R package “DiffBind” using default parameters (Ross-Innes et al., 2012; Stark, 2011).

For the analysis of RNA-seq data, sequencing reads mapping to rRNAs were filtered out using Sortmerna (Kopylova et al., 2012) and the remaining reads were trimmed of the adapter using Trimmomatic (Bolger et al., 2014). Alignment to the *A. thaliana* genome was performed with STAR (Dobin et al., 2013) and reads count with HTSeqCount (Anders et al., 2015). Differential expression analysis was performed using DESeq2 with default parameters (Love et al., 2014).

### Electrophoretic Mobility Shift Assay (EMSA)

Coding sequences of both the wild-type version as well as the phosphomimic variant (T282) of *FD* and its C-terminal domain (*FD-C*, amino acids: 203-285), *14-3-3n* (At3g02520; *GRF7*), *FT*, and *TFL1* were amplified by PCR to generate N-terminal 6X-His-tag CDS which were cloned into pETM-11 expression vector by restriction. All plasmids were transformed into *Escherichia coli* strain Rosetta plysS and proteins were induced with 1mM IPTG at 37°C over-night. Cell lysis was performed by sonication and proteins were purified using His60 columns (Clontech) and eluted in 50 mM of sodium phosphate buffer pH 8.0, 300 mM NaCl, 300 mM Imidazole. EMSA was performed using 5’-Cy3-labeled, double-stranded oligos of 30 bp covering the G-box contained in the *SEP3* promoter as a probe (Eurofins). For probe synthesis, single strand oligos were annealed in annealing buffer (10 mM Tris pH 8.0, 50 mM NaCl, 1 mM EDTA pH 8.0). Binding reactions were carried out in buffer containing 10 mM Tris pH 8.0, 50 mM NaCl, 10 µM ZnSO4, 50 mM KCl, 2.5% glycerol, 0.05% NP-40 in a total volume of 20 µl. The binding reaction was kept in dark at room temperature for 20 minutes and then loaded in native 8% polyacrylamide gel and run in 0.5X TBE at 4°C in dark. Results were visualized using a Typhoon imaging system.

## Supporting information

## AUTHOR CONTRIBUTIONS

S.C., L.Y., and M.S. designed the experiments. L.Y. established some of the FD:GFP reporter lines and performed initial ChIP (-seq) and flowering time analyses. M.N. cloned phosphomic and non-phosphorable versions of FD and analyzed their effect on flowering time. S.C. performed the EMSA studies, flowering times analysis and carried out and analyzed the ChIP-seq and RNA-seq experiments. S.C. and M.S. wrote the manuscript with input from all authors.

## ACKNOWLEDGEMENTS

We thank Diana Saez for help in isolating homozygous *myc2* and *afr1* mutants, the Protein Expertise Platform (PEP) at the Chemical Biological Center (KBC) at Umeå University for help with protein purification, and Nicolas Delhomme from the UPSC Bioinformatics Facility for assistance with submission of sequencing data. We acknowledge funding to the UPSC through grants from VINNOVA and The Knut and Alice Wallenberg Foundation. S.C. was supported through a Humboldt Foundation long-term postdoctoral fellowship. L.Y. acknowledges funding from the European Research Council (ERC) under the European Union’s Horizon 2020 research and innovation programme (grant agreement 679056) and the UK Biological and Biotechnology Research Council (BBSRC) via grant BB/P013511/1 to the John Innes Centre. Supported through the Sonderforschungsbereich 1101 (Collaborative Research Centre 1101), project grant SFB1101/1-B04, and a research project grant from the Knut and Alice Wallenberg Foundation (2016.0025) to M.S.

## SUPPLEMENTAL MATERIALS

### Supplemental figures

**Figure S1.** ChIP-seq summary statistics for the different biological replicates: *pSUC2::GFP:FD* in Col-0 (A, D, G, J) and *ft-10 tsf-1* mutant background (B, E, H, K), *pFD::GFP:FD* in *fd-2* mutant background (C, F, I, L).

**Figure S2.** Verification of comparability of controls used for normalization of FD (*pSUC2::GFP:FD*) ChIP-seq in WT and *ft-10 tsf-1* seedlings.

**Figure S3.** Effect of misexpression of FD on gene expression and flowering time.

**Figure S4.** Electrophoretic mobility shift assays (EMSAs) to test FD binding to the *SEP3* and *AP1* promoters.

**Figure S5.** Summary of RNA-seq results.

**Figure S6.** Expression profile of selected FD target genes.

**Figure S7.** Gene Ontology (GO) analysis on the subset of 135 direct genes of FD.

### Supplemental tables

**Table S1.** List of mutants and oligos for genotyping used in the study.

**Table S2.** List of vectors used in the study.

**Table S3.** List of oligos used for qRT-PCR in the study.

### Supplemental Data Sets

**Supplemental Data Set 1.**List of 1754 FD-bound peaks identified in seedlings expressing *pSUC2::GFP:FD* in Col-0.

**Supplemental Data Set 2.** List of 2427 FD-bound peaks identified in seedlings expressing *pSUC2::GFP:FD* in *ft-10 tsf-1*.

**Supplemental Data Set 3.** List of 1514 FD-bound peaks detected in seedlings expressing *pSUC2::GFP:FD* in either Col-0 or *ft-10 tsf-1*.

**Supplemental Data Set 4.** List of 917 peaks that were differential bound in seedlings expressing *pSUC2::GFP:FD* in either Col-0 or *ft-10 tsf-1*.

**Supplemental Data Set 5.** List of 595 shared FD-bound peaks in apices *pFD::GFP:FD fd-2* rescue line.

**Supplemental Data Set 6.** List of differentially expressed genes.

**Supplemental Data Set 7.** List of 135 potential direct targets of FD.

**Supplemental Data Set 8.** List of 27 genes related to “response to hormone” category within the subset of the 135 direct target of FD.

