## Supplementary material for "Effects of FLOWERING LOCUS T on FD during the transition to flowering at the shoot apical meristem of *Arabidopsis thaliana*"

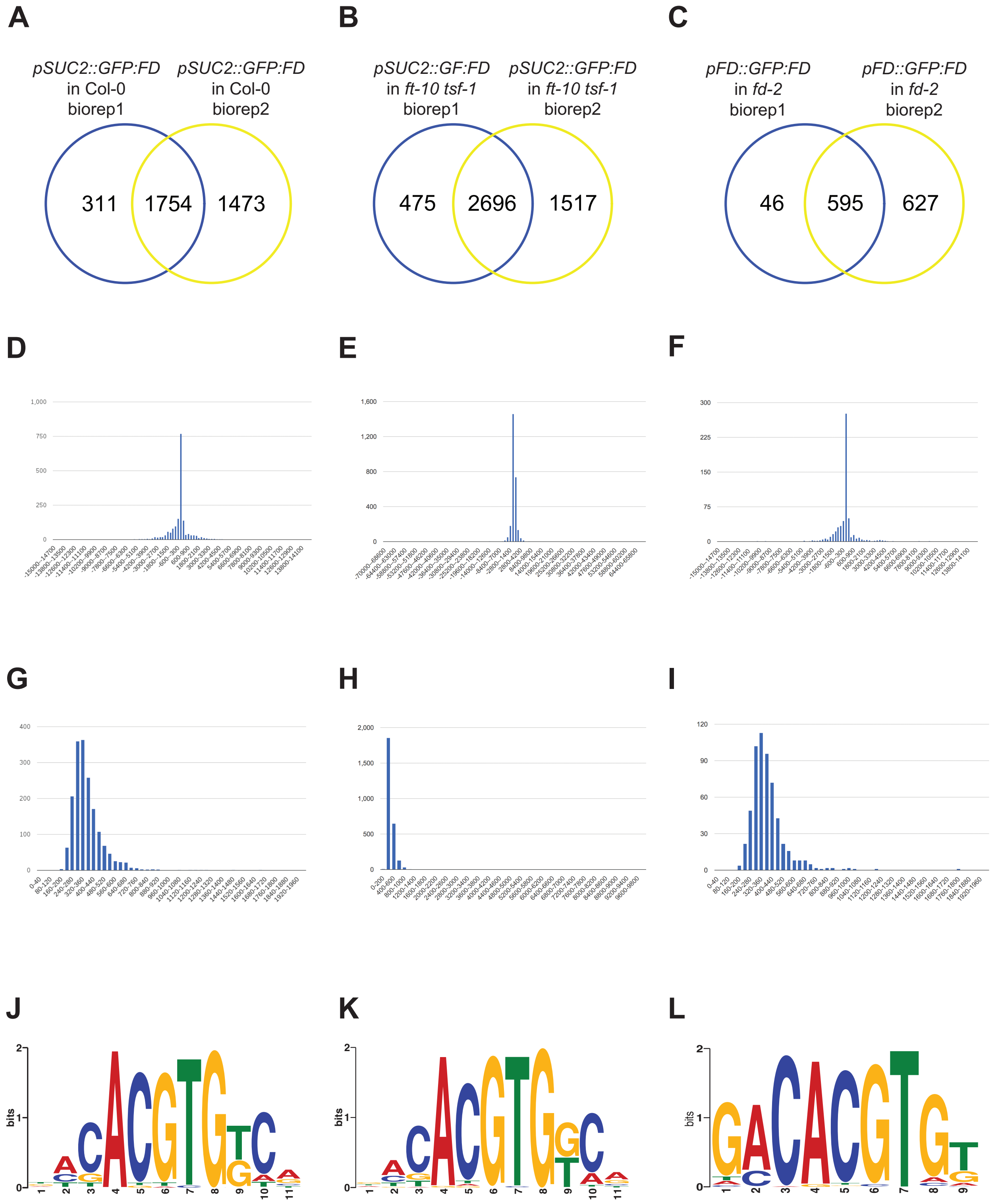


**Figure S1.** ChIP-seq summary statistics for the different biological replicates: *pSUC2::GFP:FD* in Col-0 (A, D, G, J) and *ft-10 tsf-1* mutant background (B, E, H, K), *pFD::GFP:FD* in *fd-2* mutant background (C, F, I, L).

**(A-C)** 2-set venn diagram showing the overlap of FD-bound peaks between two biological replicates.

**(D-F)** Distribution of the distance to the nearest TSS for the shared peaks.

**(G-I)** Distribution of the width of the peak for the shared peaks.

**(J-L)** Nucleotide logo of the predicted FD binding site based on peaks regions shared between biological replicates.

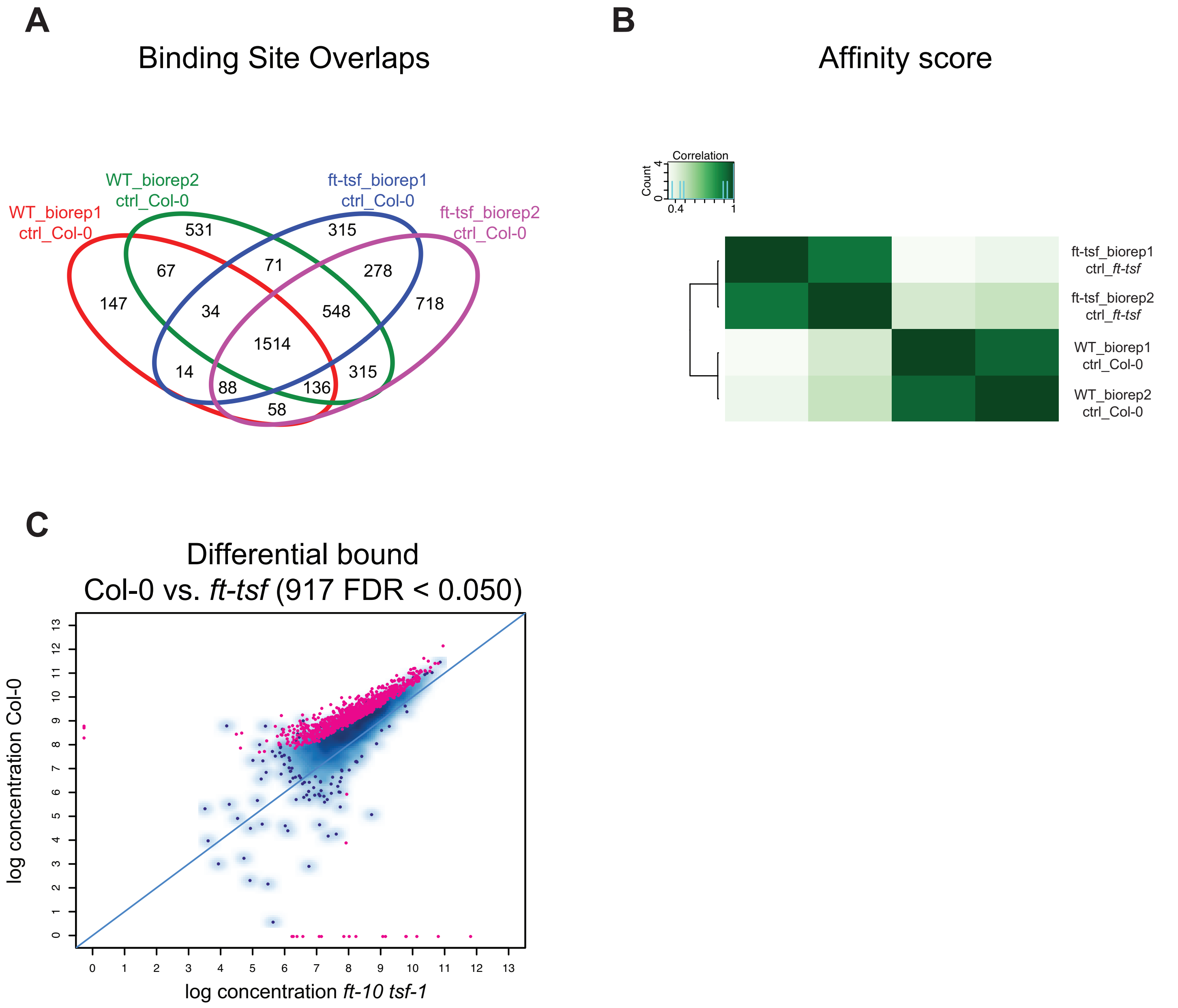


**Figure S2.** Verification of comparability of controls used for normalization of FD (*pSUC2::GFP:FD*) ChIP-seq in Col-0 and *ft-10 tsf-1* seedlings.

To test the robustness of results reported in Fig. 1B-E and to confirm that the usage of two different backgrounds as controls did not introduce any undue bias in peaks calling, we used *pSUC2::GFP:NLS* in Col-0 also for normalization of the experiments conducted in *ft-10 tsf-1*.

**(A)** 4-set venn diagram of the overlapping of the four biological replicates using only the controls in Col-0. Numbers are very similar to the ones reported in Fig. 1B.

**(B)** Binding matrix based on affinity scores confirms that biological replicates still cluster by genotype.

**(C)** A total of 917 peaks were found as differentially bound (FDR < 0.05) between WT and *ft-10 tsf-1* using *pSUC2::GFP:NLS* in Col-0 as the unique control, which is very similar to 885 reported in Fig. 1E.

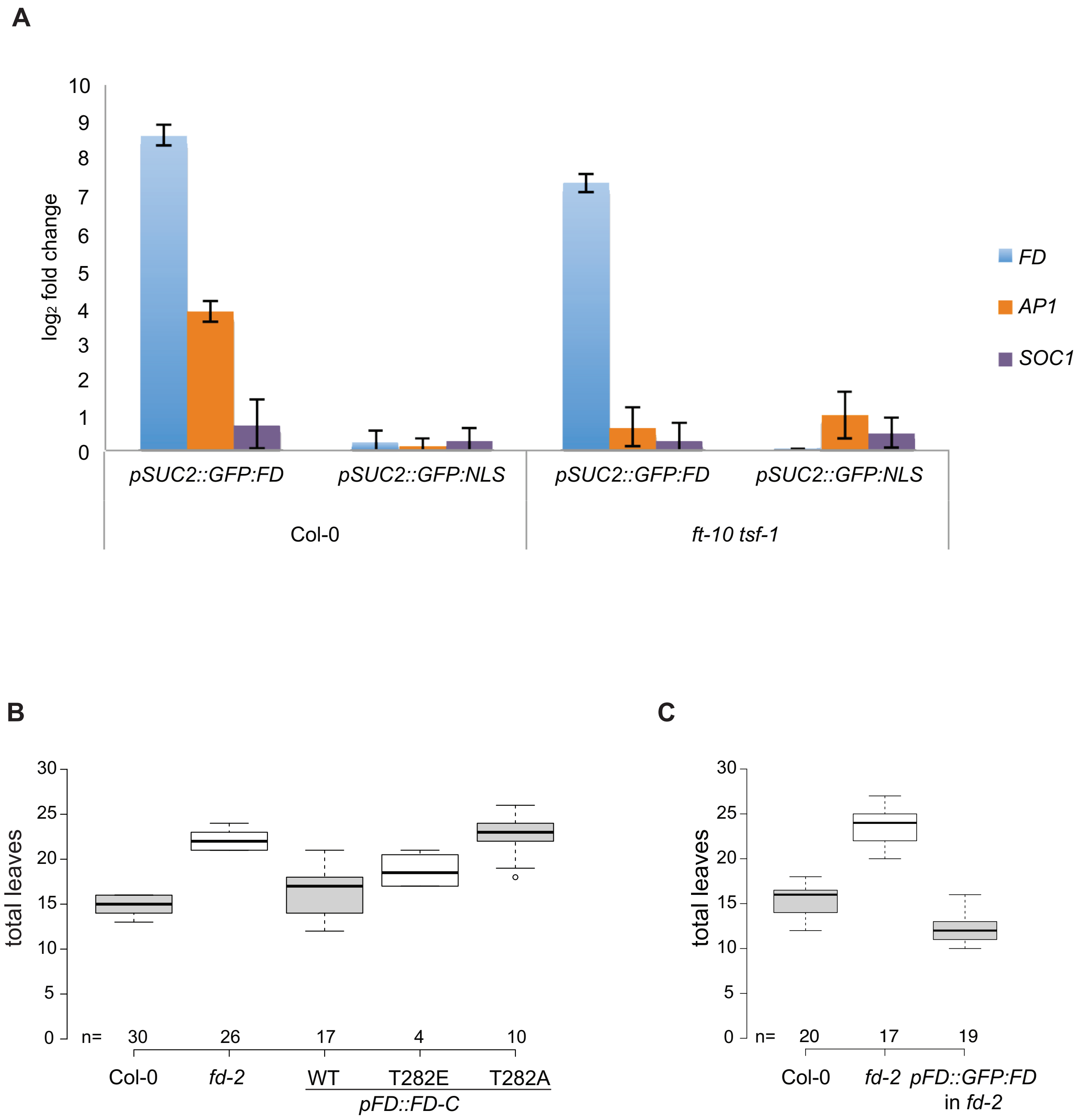


**Figure S3.** Effect of misexpression of FD on gene expression and flowering time.

**(A)** Expression analysis of *FD*, *AP1* and *SOC1* in *pSUC2::GFP:FD* and *pSUC2::GFP:NLS* in Col-0 and *ft-10 tsf-1* mutant. Leaves were collected from 16 days old seedlings carrying *pSUC2::GFP:FD* and *pSUC2::GFP:NLS* both in Col-0 and *ft-10 tsf-1* mutant grown under LD at 23ºC. *FD* is strongly induced in leaves only when expressed under SUC2 promoter. The strong induction of *FD* in leaves induce only the expression of *AP1,* but not of *SOC1*, in Col-0 compared to *ft-10 tsf-1* mutant.

**(B)** Flowering time of *fd-2* expressing the C-terminal fragment of FD under control of the pFD promoter. The unaltered version (WT), the phosphomic version (T282E), and the non-phosphorable version (T282A) of the C-terminal fragment of FD (amino acids 203 – 285) were transformed in to *fd-2* and flowering time was scored in T1 plants after Basta treatment. Col-0 and *fd-2* plants were used as controls. Number of independent T1 plants is indicated.

**(C)** Complementation of *fd-2* by *pFD::GFP:FD*. Flowering time of *fd-2* mutants transformed with *pFD::GFP:FD* to demonstrate the activity of GFP:FD fusion protein. Flowering time was scored as total leaves number (B). Col-0 and *fd-2* plants were used as controls. Number of plants is indicated.

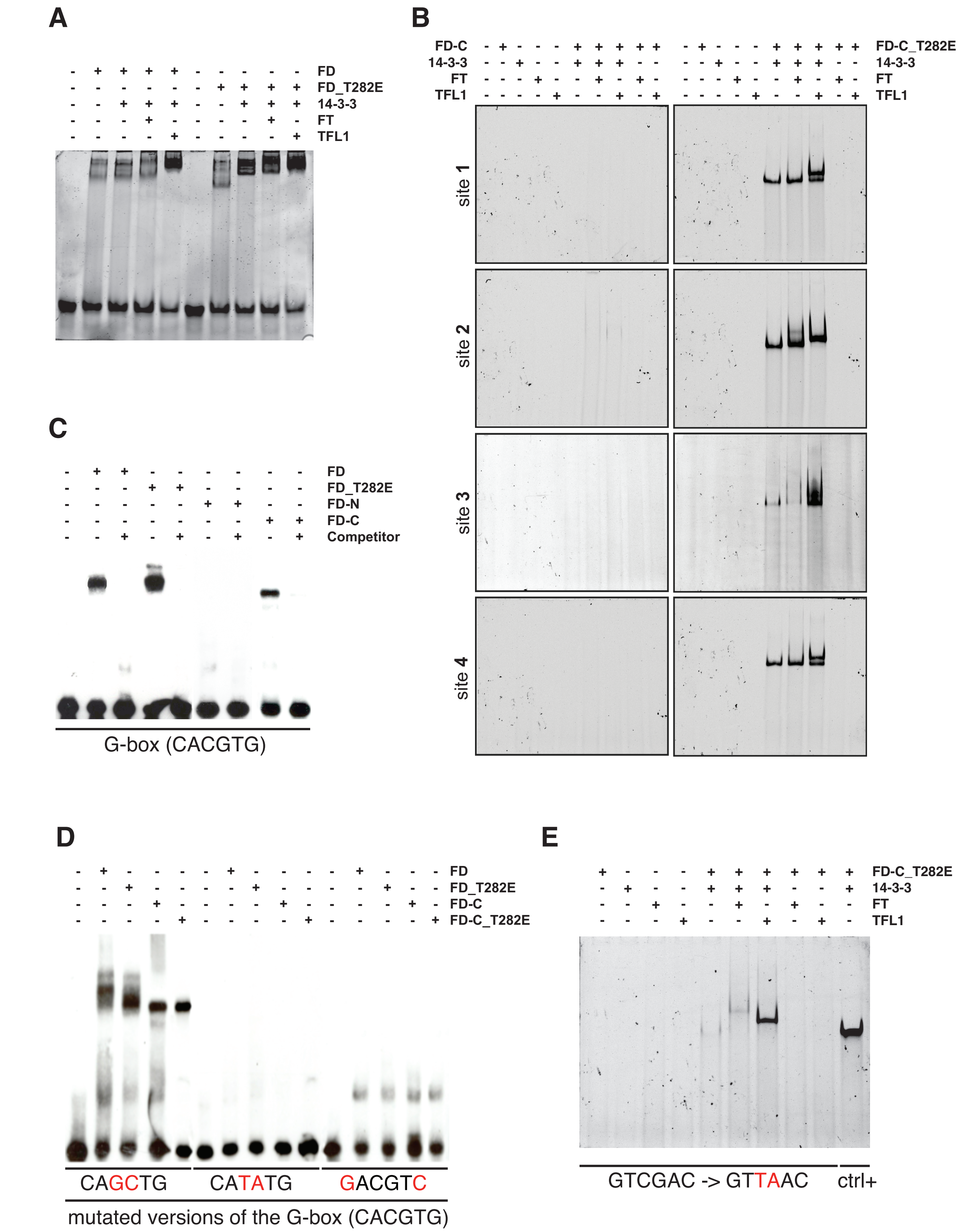


**Figure S4.** Electrophoretic mobility shift assays (EMSAs) to test FD binding to the *SEP3* and *AP1* promoters.

- **(A)** Wildtype and phosphomic (T282E) versions of full-length FD protein bind the G-box from *SEP3* promoter in the absence of interaction partners. Higher order complexes are formed in the presence of 14-3-3v and FT or TFL1, essentially confirming results obtained using a C-terminal fragment of FD (amino acids 203 – 285; Fig. 2).
- **(B)** Comparison of the wildtype and phosphomimic version of FD-C (FD-C_T282E) in combinations with 14-3-3v, FT and TFL1 using the four putative binding sites in the *AP1* promoter reported in Fig. 4B. Free probes are not visible because gels were running longer to maximize the distance between shifted probes. Only the phospho-mimic version of FD-C (FD-C_T282E) forms higher order complexes and binds DNA.
- **(C)** EMSAs to verify whether the C-terminal of FD alone (FD-C), which contains the DNA-binding domain, is sufficient to bind DNA and whether the N-terminal can interact with DNA.
- **(D)** EMSAs to test the influence of different mutations of the G-box in the interaction of FD with DNA.
- **(E)** EMSA to test whether the mutation on the site 2 on AP1 promoter is sufficient to abolish the interaction of FD with DNA reported in Fig. 4B. In “ctrl+” the probe containing the G-box from SEP3 was used.

EMSA reported in panels C and D were preliminar experiments in order to set the assay and were done with Chemiluminescent EMSA kit (ThermoFisher Scientific cat. 20148) according to the manufacters instructions. We then shifted to fluorescent labeled probes (see methods in the main text) because it is a faster, cheaper and more accurate method. Images reported in these panels come from different gels and were merged together for a better representation.

A

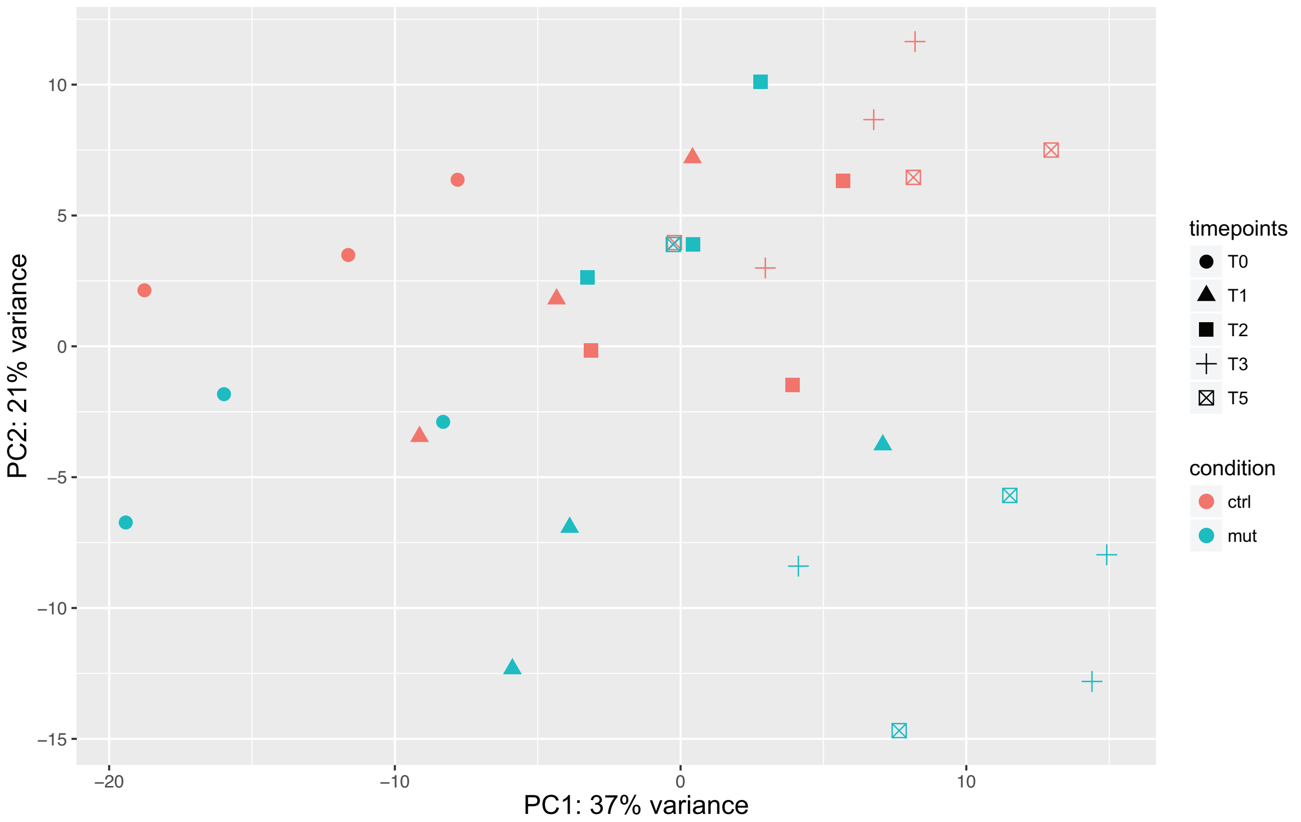


B

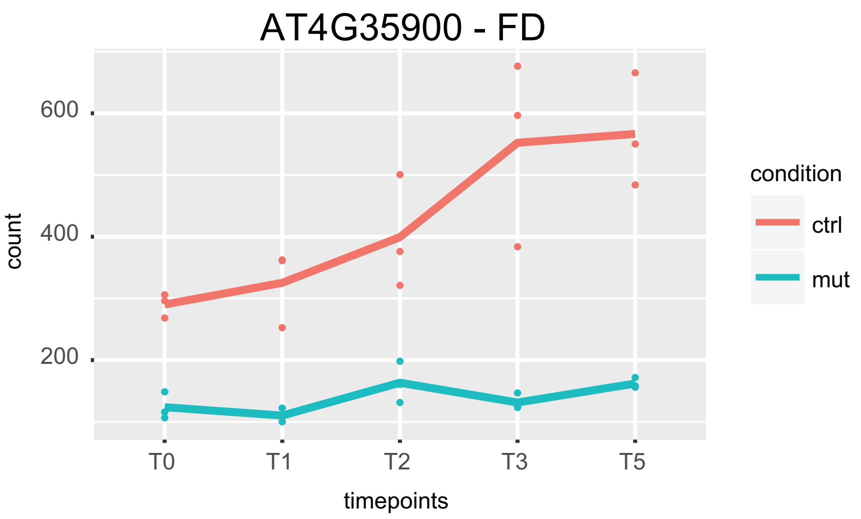


**Figure S5.** Summary of RNA-seq results.

**(A)** PCA analysis of 30 RNA-seq samples. 37% of the variance in the data set is explained by the different time points and 21% by the genotypes. Genotypes become visibly separated in time points T3 and T5. Red marks control samples (ctrl.; *pFD::GFP:FD fd-2*) and blue is used for *fd-2* mutant.

**(B)** Expression profile of FD in control and *fd-2* mutants. The expression of FD increases after T2 in agreement with the results showed in (A). Red dots indicate gene expression in control samples (*pFD::GFP:FD fd-2*), blue dots indicate gene expression in *fd-2*. Mean expression in control and *fd-2* is indicated by red and blue lines, respectively.

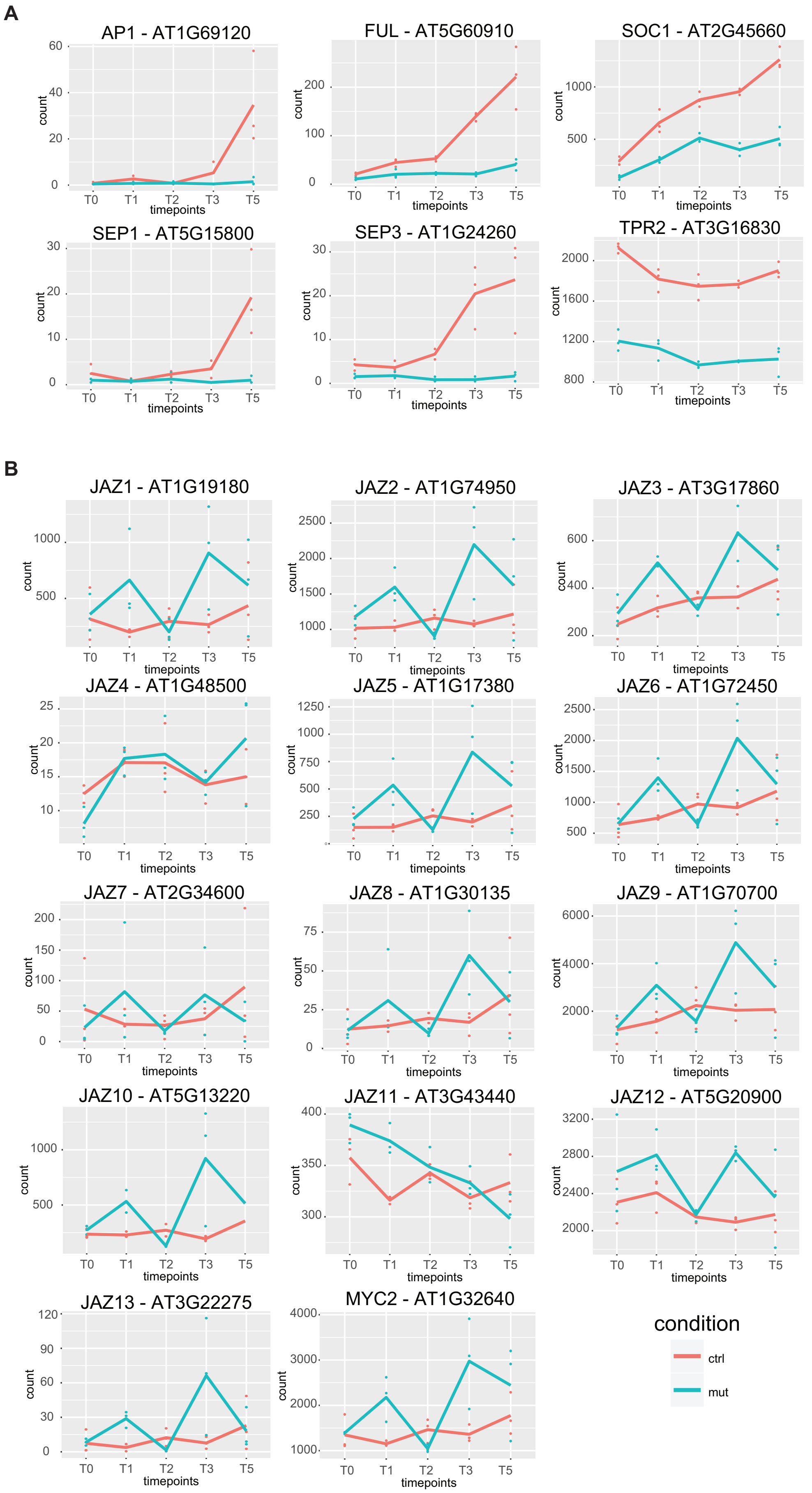


**Figure S6.** Expression profile of selected FD target genes.

**(A)** Expression profiles of *FD*, *AP1*, *FUL*, *SOC1*, *SEP3*, *SEP1*, and *TPR2* are reported.

**(B)** Expression profiles of 13 *JAZ* genes and *MYC2* at the shoot apical meristem for five time points during floral transition are shown.

Red dots indicate gene expression in control samples (*pFD::GFP:FD fd-2*), blue dots indicate gene expression in *fd-2*. Mean expression in control and *fd-2* is indicated by red and blue lines, respectively.

**A**

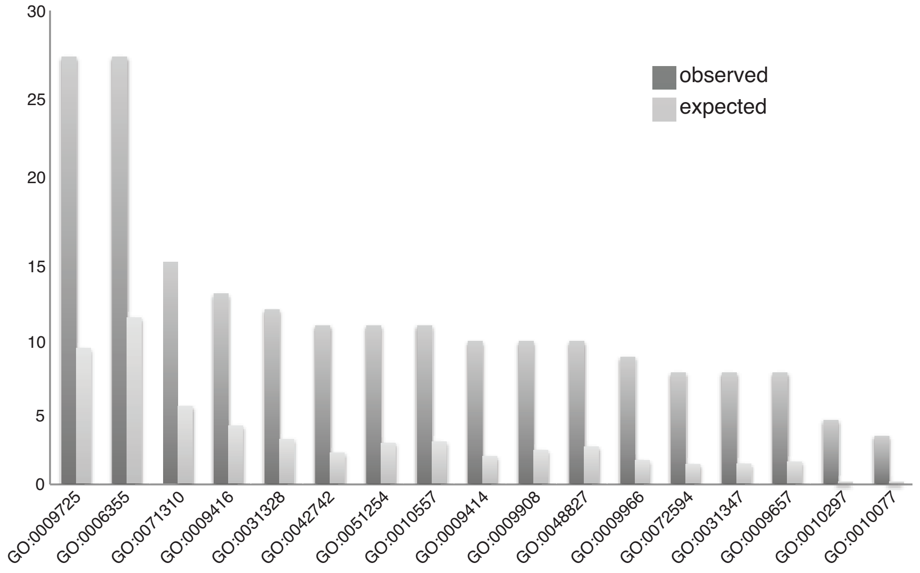


GO:0009725 response to hormone
GO:0006355 regulation of transcription, DNA-templated
GO:0071310 cellular response to organic substance
GO:0009416 response to light stimulus
GO:0031328 positive regulation of cellular biosynthetic process
GO:0042742 defense response to bacterium
GO:0051254 positive regulation of RNA metabolic process
GO:0010557 positive regulation of macromolecule biosynthetic process
GO:0009414 response to water deprivation
GO:0009908 flower development
GO:0048827 phyllome development
GO:0009966 regulation of signal transduction
GO:0072594 establishment of protein localization to organelle
GO:0031347 regulation of defense response
GO:0009657 plastid organization
GO:0010207 photosystem II assembly
GO:0010077 maintenance of inflorescence meristem identity

**B**

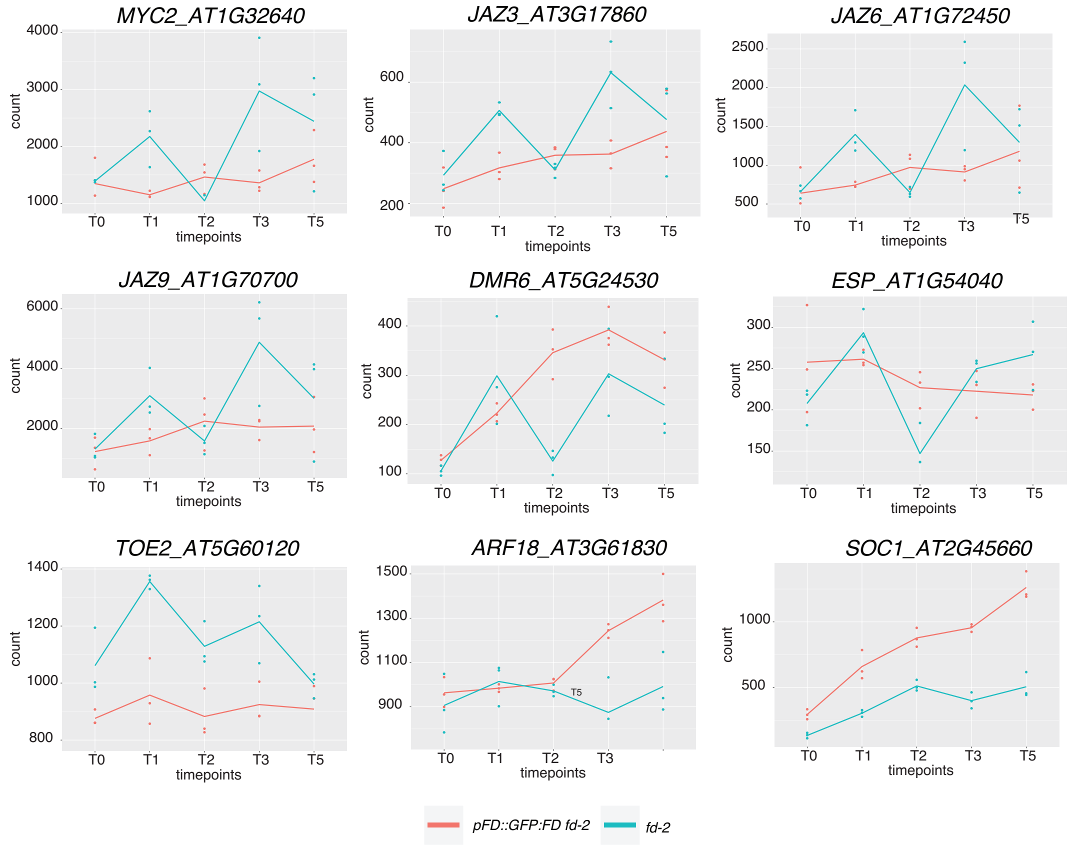


**Figure S7.** Gene Ontology (GO) analysis on the subset of 135 direct genes of FD.

**(A)** Significantly enriched GO categories (FDR < 0.05).

**(B)** Genes with the peculiar expression profile related to pathogen resistance and jasmonate pathway and *ARF18*, gene related to auxin, which share the expression profile with *SOC1*, gene related to gibberellin signaling and important flowering-related gene.

**Table S1.** List of mutants and oligos for genotyping used in the study.

| line | locus | SALK | oligos for screening the mutant | | |
|---|---|---|---|---|---|
| *fd-2* | AT4G35900 | SALK_013288 | WT | Forward | GAAAATAGAAAGTGAGATAAAACC |
| | | | | Reverse | TGGAAAAGAGAACAGAAGTGAACC |
| | | | mut | Forward | ATTTTGCCGATTTCGGAAC |
| | | | | Reverse | TTCCAAACTTCTTCCATGGTG |
| *ft-10* | AT1G65480 | GABI_290E08 | WT | Forward | ATATTGATGAATCTCTGTTGTGG |
| | | | | Reverse | AGGGTTGCTAGGACTTGGAACA |
| | | | mut | Forward | CCCATTTGGACGTGAATGTAGACAC |
| | | | | Reverse | AGGGTTGCTAGGACTTGGAACA |
| *tsf-1* | AT4G20370 | SALK_087522 | WT | Forward | CGGTTAACTTGATTTTGTTTCG |
| | | | | Reverse | ACGTGGACTCTCGTAGCACAC |
| | | | mut | Forward | ATTTTGCCGATTTCGGAAC |
| | | | | Reverse | ACGTGGACTCTCGTAGCACAC |
| *myc2* | AT1G32640 | SALK_017005C | WT | Forward | CCTACGCTATATTCTGGCAACC |
| | | | | Reverse | AGTGGCTCTTCTCTACCGTTTG |
| | | | mut | Forward | ATTTTGCCGATTTCGGAAC |
| | | | | Reverse | AGTGGCTCTTCTCTACCGTTTG |
| *afr1* | AT1G75060 | SALK_026979C | WT | Forward | TTGGCTTAAGAATCACTCCATG |
| | | | | Reverse | AAAGCGAAGTTGATCTTTGCTC |
| | | | mut | Forward | ATTTTGCCGATTTCGGAAC |
| | | | | Reverse | AAAGCGAAGTTGATCTTTGCTC |

**Table S2.** List of of vectors used in the study.

| Construct | Description | Resistance |
|---|---|---|
| pLY-33 | *pGREEN - pSUC2::GFP:FD* | Spectinomycin |
| pLY-100 | *pGREEN - pSUC2::GFP:NLS* | Spectinomycin |
| pLY-46 | *pGREEN - pFD::GFP:FD* | Spectinomycin |
| pJM-54 | *pGREEN - pFD::FD* | Spectinomycin |
| pMH-52 | *pGREEN - pFD::FD-T282A* | Spectinomycin |
| pMH-54 | *pGREEN - pFD::FD-T282E* | Spectinomycin |
| pMH-60 | *pGREEN - pFD::FD-S281E* | Spectinomycin |
| pMH-58 | *pGREEN - pFD::FD-S281E/T282E* | Spectinomycin |
| pSC-215 | *pGREEN - pFD::C-FD* | Spectinomycin |
| pSC-216 | *pGREEN - pFD::C-FD-T282A* | Spectinomycin |
| pSC-217 | *pGREEN - pFD::C-FD-T282E* | Spectinomycin |
| pSC-098 | *pET-M11 - 6X-His-FD* | Kanamycin |
| pSC-099 | *pET-M11 - 6X-His-FD-T282E* | Kanamycin |
| pSC-100 | *pET-M11 - 6X-His-14-3-3(Nu)* | Kanamycin |
| pSC-101 | *pET-M11 - 6X-His-C-FD* | Kanamycin |
| pSC-102 | *pET-M11 - 6X-His-C-FD-T282E* | Kanamycin |
| pSC-130 | *pET-M11 - 6X-His-FT* | Kanamycin |
| pSC-166 | *pET-M11 - 6X-His-TFL1* | Kanamycin |

**Table S3.** List of of oligos used for qRT-PCR in the study.

| gene | locus | oligos for qRT-PCR | |
|---|---|---|---|
| FD | AT4G35900 | Forward | GCAAGACTCAAGAGACAACAAG |
| | | Reverse | CAAAATGGAGCTGTGGAAGAC |
| SOC1 | AT2G45660 | Forward | AAACGAGAAGCTCTCTGAAAAG |
| | | Reverse | AAGAACAAGGTAACCCAATGAAC |
| AP1 | AT1G69120 | Forward | CACCAAATCCAGCATCCTTAC |
| | | Reverse | AGTTCGAGATCATTCCTCCTC |
| AS1 | AT2G37630 | Forward | AGAGAGCAGAGAACGGTCCAGG |
| | | Reverse | TCGGTGCCCTTCCTCCAACTCT |
| SEP1 | AT5G15800 | Forward | ATGATTGGTGTGAGAAGTCATCATATG |
| | | Reverse | GATGTAACCGTTTCCCTGCTGCGCCTG |
| SEP2 | AT3G02310 | Forward | ATCAACAGAATATTGCCTATGGACATC |
| | | Reverse | GATGTAGCCGTTTCCTTGTTGGGACTG |
| SEP3 | AT1G24260 | Forward | GGGTATCAGATGCCACTCCAGCTGAAC |
| | | Reverse | AACCCAACATGTAATTATTCACACTTG |
| SEP4 | AT2G03710 | Forward | GAGAAAGTTGGAGGACAGTGATGC |
| | | Reverse | GCTCATGCCTTGTTGCTGTTGT |
| FRI-like 4a | AT3G22440 | Forward | GGAGGACTCTAGCAATACTGGCCG |
| | | Reverse | AGCTGATCCAACCGTTTCTTGAGG |
| FRI-like 4b | AT4G14900 | Forward | CCAACAATTCTGGCCGATCTGC |
| | | Reverse | CGGGAATCACGGCTGGTTTTCT |
| MYC2 | AT1G32640 | Forward | CGGTGGGGATGGAGATTGAAGTGA |
| | | Reverse | TCAACGCCGACATCAACCTCGC |
| AFR1 | AT1G75060 | Forward | TGCTCTTCCGAATCCCACAAAG |
| | | Reverse | TGCAGCCTGAACGAATCCCACA |
